# Lipid hydroperoxides promote sarcopenia through carbonyl stress

**DOI:** 10.1101/2021.12.17.473200

**Authors:** Hiroaki Eshima, Justin L. Shahtout, Piyarat Siripoksup, MacKenzie J. Pearson, Ziad S. Mahmassani, Patrick J. Ferrara, Alexis W. Lyons, J. Alan Maschek, Alek D. Peterlin, Anthony R. P. Verkerke, Jordan M. Johnson, Anahy Salcedo, Jonathan J. Petrocelli, Edwin R. Miranda, Ethan J. Anderson, Sihem Boudina, Qitao Ran, James E. Cox, Micah J. Drummond, Katsuhiko Funai

## Abstract

Reactive oxygen species (ROS) accumulation is a cardinal feature of skeletal muscle atrophy. ROS refers to a collection of radical molecules whose cellular signals are vast, and it is unclear which downstream consequences of ROS are responsible for the loss of muscle mass and strength. Here we show that lipid hydroperoxides (LOOH) are increased with age and disuse, and the accumulation of LOOH by deletion of glutathione peroxidase 4 (GPx4) is sufficient to augment muscle atrophy. LOOH promoted atrophy in a lysosomal-dependent, proteasomal-independent manner. In young and old mice, genetic and pharmacologic neutralization of LOOH or their secondary reactive lipid aldehydes robustly prevented muscle atrophy and weakness, indicating that LOOH-derived carbonyl stress mediate age- and disuse-induced muscle dysfunction. Our findings provide novel insights for the role of LOOH in sarcopenia including a therapeutic implication by pharmacologic suppression.

## Introduction

Loss of muscle mass and function with age is detrimental to health and quality of life [1, 2]. Sarcopenia, muscle atrophy and weakness with aging, is due to a combination of inactivity, injury, surgery, and biological consequences of aging [3, 4]. A pharmacologic therapy for muscle loss does not exist, and current diet or exercise therapeutic approaches are often ineffective or unfeasible. Oxidative stress has been implicated in muscle atrophy by accelerating proteolysis [5, 6], but the exact mechanism by which reactive oxygen species (ROS) contributes to the decrease in muscle mass and strength is not well understood.

Lipid hydroperoxide (LOOH) is a class of ROS molecules that has been implicated in cell damage, particularly as a trigger to induce ferroptosis, a non-apoptotic form of regulated cell death [7, 8]. Lipid peroxidation is initiated by prooxidants such as hydroxyl radicals attacking the carbon-carbon double bond in fatty acids, particularly the polyunsaturated fatty acids (PUFAs) containing phospholipids [9]. Lipid radicals (L•) created by this reaction rapidly reacts with oxygen to form a lipid peroxy-radical which subsequently reacts with another lipid to produce L• and LOOH, the former propagating lipid peroxidation. LOOH is the primary product of lipid peroxidation that forms secondary reactive lipid aldehydes such as 4-hydroxynonenal (4-HNE) and malondialdehyde (MDA), inducing carbonyl stress with high reactivity against biological molecules to promote cellular toxicity. The intracellular level of LOOH is endogenously suppressed by glutathione peroxidase 4 (GPx4) that catalyzes the reaction by which LOOH is reduced to its nonreactive hydroxyl metabolite [10].

Despite the evidence for the role of LOOH-mediated cell damage and cell death, the biological consequence of LOOH accumulation in skeletal muscle is not well understood [11, 12]. Below we provide evidence that LOOH mediates the loss of muscle mass and function associated with sarcopenia. An increase in muscle LOOH was a common feature with aging and disuse, and accumulation of LOOH in vitro and in vivo augmented muscle atrophy. We further show that genetic or pharmacologic suppression of LOOH and their reactive lipid aldehydes is sufficient to prevent disuse-induced muscle atrophy in young and old mice.

## Results

We first evaluated the changes in skeletal muscle LOOH with aging. In humans and in mice, aging promoted a reduction in the expression of GPx4 in skeletal muscle (Fig. 1A&B). To examine the changes in skeletal muscle LOOH landscape with age, we performed a comprehensive oxidolipidomic analysis in muscle samples from young (4 months) and old (20 months) mice (Fig. 1C&D). We detected over 300 species of oxidized lipids with an effect distribution that was highly class-dependent. Among these, age had the most robust effect on oxidized phosphatidylethanolamine (Fig. 1C, red), a class of lipids that have been implicated as a potential lipid signal to induce ferroptosis [13]. Among the top ten oxidized lipid species whose abundance was most robustly increased with age, six of them were oxidized phosphatidylethanolamine (Figure 1D), and they were substantially more highly abundant compared to other oxidized lipids. LOOH can be indirectly assessed by quantifying lipid aldehyde adducts such as 4-hydroxynonenal (4-HNE) and malondialdehyde (MDA). We confirmed increased muscle 4-HNE and MDA with age (Fig. 1E-G).

**Fig. 1.**
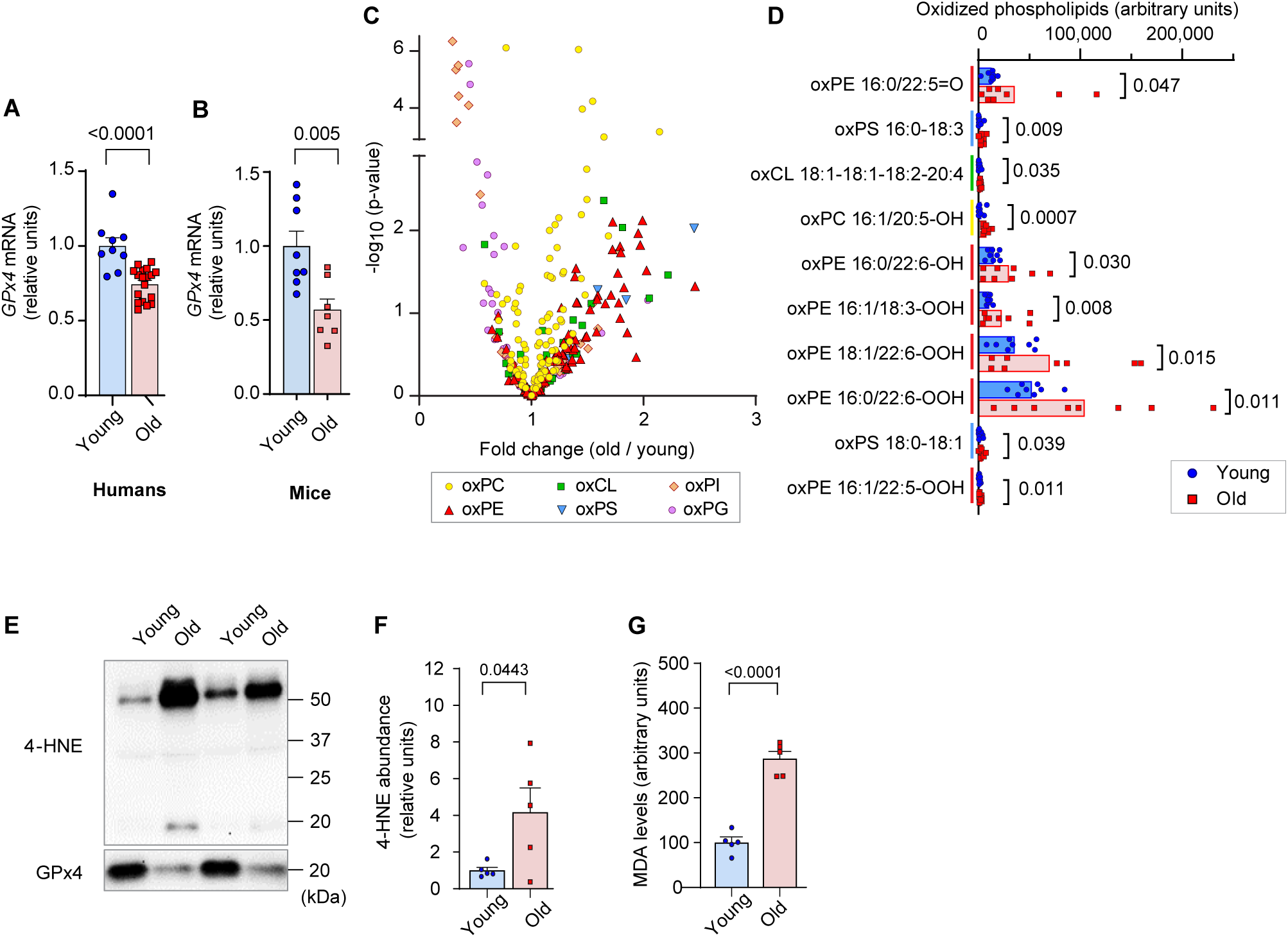
LOOH increases with age in skeletal muscle. **(A,B)** GPx4 mRNA levels in skeletal muscle biopsy samples from young and old humans (*n =* 9 for young, *n =* 18 for old) **(A)** or skeletal muscles from young and old mice (*n =* 8 for young, *n =* 7 for old) **(B)**. **(C,D)** Oxidized phospholipid content in skeletal muscle from young and old mice (*n =* 8 per group). (**E**,**F**,**G)** Immunoblotting **(E)** and quantification **(F)** of 4-HNE proteins (*n =* 5 per group) and MDA levels **(G)** (*n =* 3 per group). Data are shown as the mean ± SEM. Statistical analyses in **(A**,**B**,**C**,**D,F)** and (**G)** were performed with an unpaired two-tailed t-test.

Disuse promoted by inactivity, injury, or surgery is a major contributor to age-associated decline in muscle mass and function. Disuse also promotes skeletal muscle atrophy that is likely contributed by ROS [5]. To model disuse atrophy, mice underwent a hindlimb unloading (HU) procedure as previously described [14, 15] (Supplemental Fig. S1A-J). As expected, HU induced muscle atrophy and weakness (Supplemental Fig. S1D&E) concomitant to reduction in body and lean mass (Supplemental Fig. S1B&C). Disuse robustly elevated muscle LOOH levels (Supplemental Fig. S1F&G) without significant changes in mitochondrial bioenergetics (Supplemental Fig. S1H-J). An increase in muscle LOOH preceded atrophy (Supplemental Fig. S1D&F), consistent with the notion that LOOH may trigger mechanisms to promote loss of muscle mass.

Next, we tested our hypothesis that LOOH contributes to muscle atrophy using C2C12 myotubes (Fig. 2A). Lentivirus-mediated knockdown (KD) of GPx4 increased LOOH and markers of ferroptosis concomitant with a decrease in myotube diameter (Fig. 2B-I). We also recapitulated these findings with erastin (a system X_c_^−^ inhibitor that suppresses glutathione synthesis) (Fig. 2B&C and Supplemental Fig. S2A-E) and RSL3 (GPx4 inhibitor) (Fig. 2B&C and Supplemental Fig. S2F-I), commonly used acute pharmacological interventions to elevate intracellular LOOH. These data support the idea that LOOH reduces myotube size in a cell-autonomous manner.

**Fig. 2.**
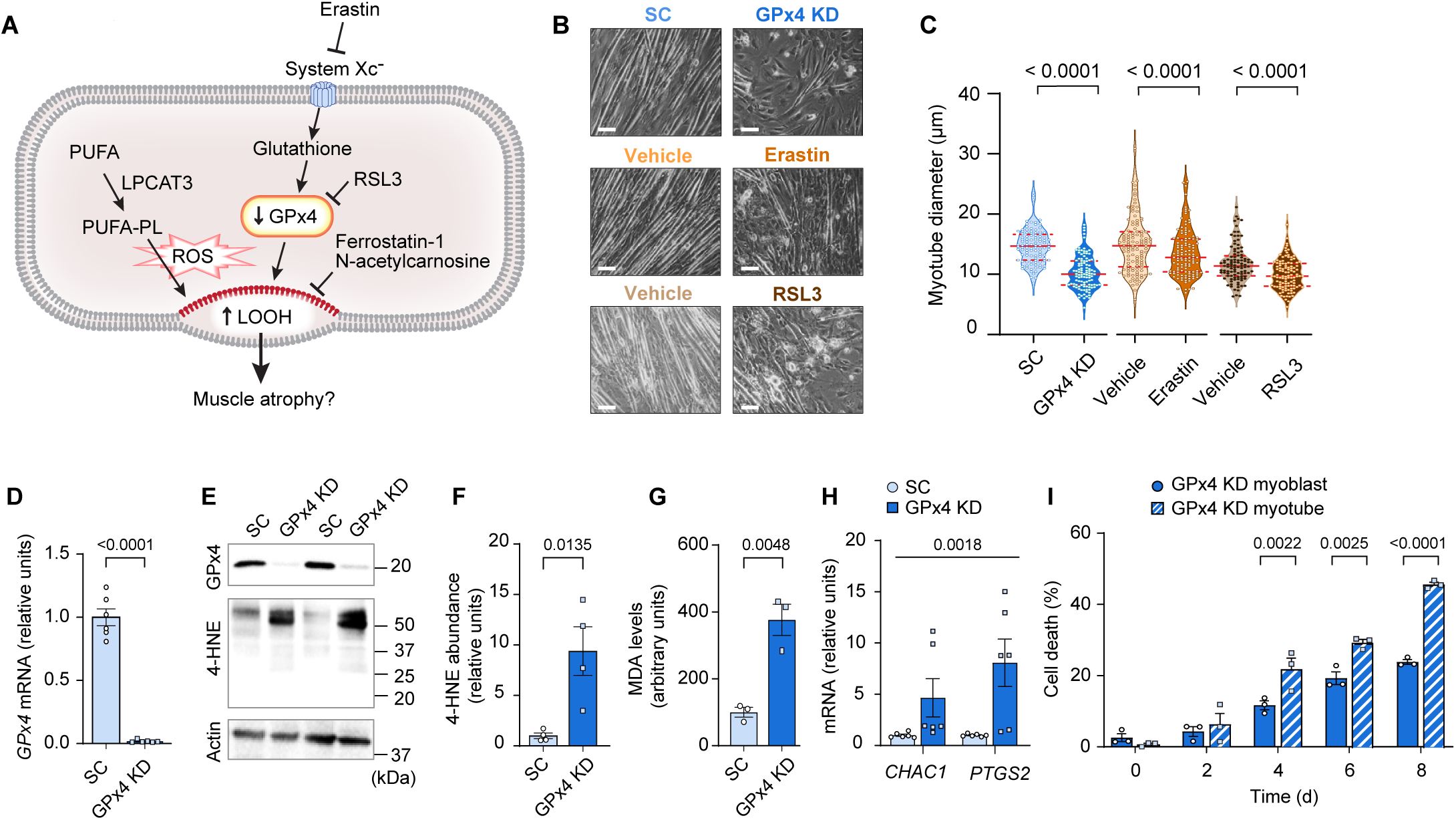
Elevated LOOH is sufficient to promote atrophy in cultured myotubes. **(A)** A schematic of how pathways that regulate LOOH may promote muscle atrophy. PUFA: phospholipids containing polyunsaturated fatty acids. **(B,C)** Representative images **(B)** and quantification **(C)** of myotube diameter (*n =* 104 for scrambled: SC, *n =* 107 for GPx4 KD, *n =* 117 for Vehicle, *n =* 120 for erastin, *n =* 104 for Vehicle, *n =* 110 for RSL3). Scale bar, 100 μm. **(D**) GPx4 mRNA levels in C2C12 myotubes with or without GPx4 knockdown (GPx4 KD) (*n =* 6 per group). **(E,F,G)** Immunoblotting of 4-HNE, GPx4, and actin **(E)**, quantification of 4-HNE **(F)** proteins and MDA levels **(G)** (*n =* 3 per group). **(H)** mRNA levels for CHAC1 and PTGS2, markers of ferroptosis (*n =* 6 per group). **(I)** cell death levels in GPx4 KD myoblast or myotubes (*n =* 3 independent repeats). Data are shown as the mean ± SEM. Statistical analyses in **(C,D,F,G)** and **(I)** were performed with an unpaired two-tailed t-test. Statistical analyses in **(H)** were performed with a two-way ANOVA and Tukey’s multiple comparison test.

We then translated these findings in vivo with global heterozygous GPx4 knockout mice (GPx4^+/−^). Germline deletion of GPx4 is embryonically lethal [16], but GPx4^+/−^ mice appear normal and do not have an observable muscle phenotype at baseline [10, 17]. We studied 4 months (young) and 20 months (old) GPx4^+/−^ and wildtype littermates with or without HU (Supplemental Fig. S3A-D). In young mice, GPx4 haploinsufficiency augmented the loss in muscle mass induced by HU (Fig. 3A and Supplemental Fig. S3E). However, muscle masses between old GPx4^+/−^ and wildtype mice were not different. We interpret these findings to mean that disuse in old mice promotes an increase in LOOH that has already reached a maximally effective threshold with age such that GPx4 deletion had no further effect. In support of this, we saw no differences in 4-HNE or MDA levels between old GPx4^+/−^ and wildtype mice (Fig. 3B and Supplemental Fig. S3F&G). GPx4 haploinsufficiency did not alter force-generating capacity (Fig. 3C and Supplemental Fig. S3H-J).

**Fig. 3.**
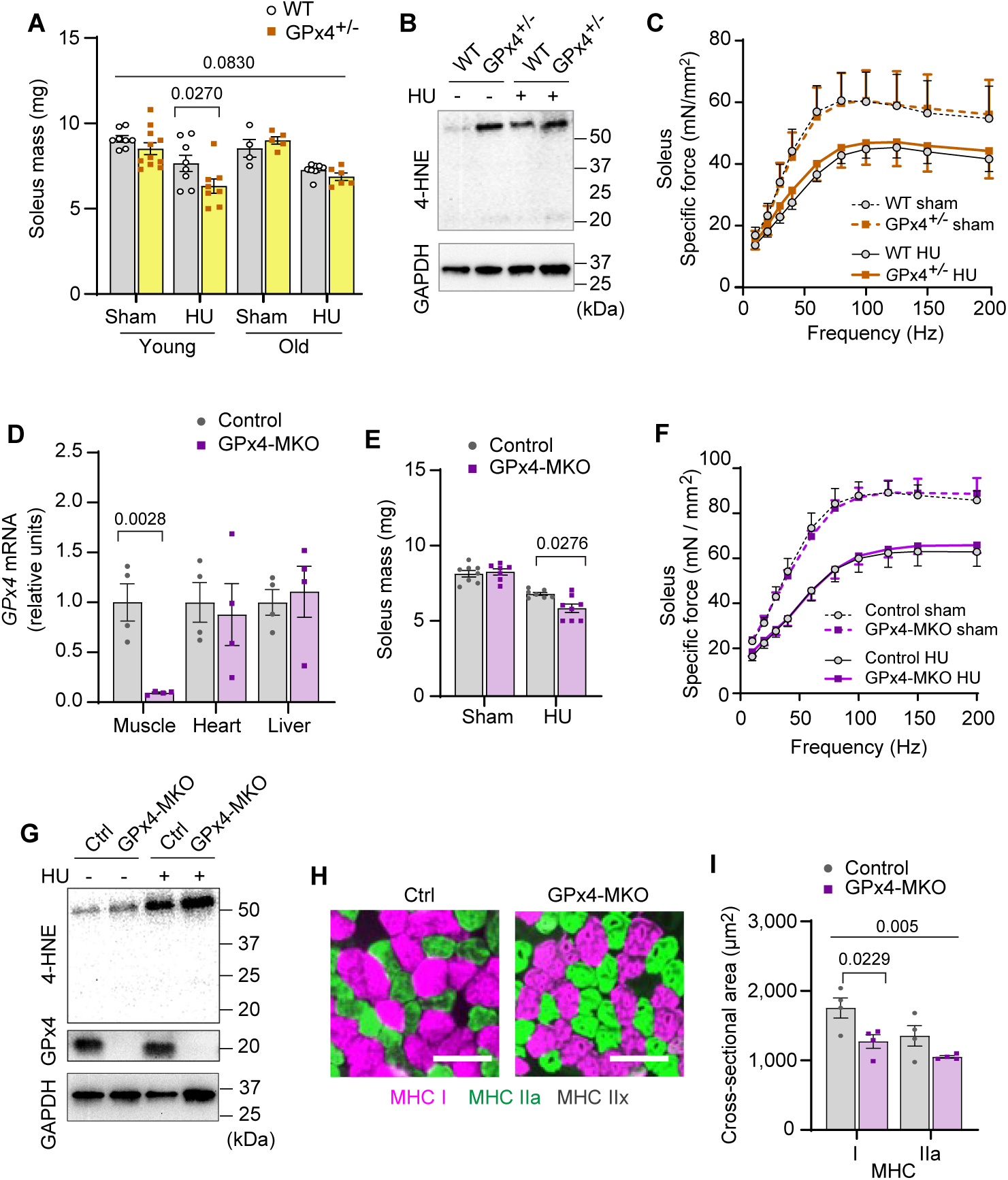
Elevated LOOH is sufficient to augment disuse-induced muscle atrophy in young and old mice. **(A)** Soleus muscle mass from young or old WT or GPx4^+/−^ mice with or without HU (*n =* 8-11 per young group, *n =* 4-8 per old mice group). **(B)** Immunoblotting of 4-HNE from old WT or GPx4^+/−^ mice. **(C)** Force-frequency curve from old WT or GPx4^+/−^ mice (*n =* 4-7 per group). **(D)** mRNA levels of GPx4 from young control or GPx4-MKO mice (*n =* 4 per group). **(E,F)** Soleus muscle mass **(E)** (*n =* 7-8 per group) or force-frequency curve **(F)** from young control or GPx4-MKO mice (*n =* 4-7 per group). **(G)** Immunoblotting of 4-HNE and GPx4 from young GPx4-MKO. **(H,I)** Representative images of MHC immunofluorescence **(H)** and muscle fiber CSA by fiber type **(I)** for soleus muscles in young control or GPx4-MKO mice with HU (*n =* 4 per group). Scale bar, 100µm. Data are shown as the mean ± SEM. Statistical analyses in **(D)** were performed with an unpaired two-tailed t-test. Statistical analyses in **(A,C,E,F)** and **(I)** were performed with a two-way ANOVA and multiple comparisons were performed using Tukey’s **(C,E,F,I)** or Sidak’s **(A)** multiple comparisons tests.

Because GPx4 is expressed globally, we also studied mice with skeletal muscle-specific tamoxifen-inducible GPx4 knockout (GPx4-MKO) (Fig. 3D and Supplemental Fig. S4A) [18]. Consistent with GPx4+/− mice, GPx4-MKO mice were also more prone to developing disuse-induced skeletal muscle atrophy (Fig. 3E&F and Supplemental Fig. S4B-F) concomitant to elevated LOOH (Fig. 3G and Supplemental Fig. S4G&H), suggesting that loss of GPx4 in muscle augments atrophy in a cell-autonomous manner. Histological analyses revealed that reduced muscle mass was consistent with reduced cross-sectional area of myofibers regardless of fiber-type compositions (Fig. 3H&I and Supplemental Fig. S4I&J). These data implicate that LOOH directly reduces muscle cell size in vivo.

GPx4 primarily neutralizes LOOH but it also exhibits some activity towards other peroxides [19]. To confirm that the effects of GPx4 deletion to promote atrophy is specific to LOOH, we diminished the ability of cells to incorporate PUFAs into phospholipids by deleting lysophosphatidylcholine acyltransferase 3 (LPCAT3) [20–22]. LPCAT3 is an enzyme of Lands cycle that preferentially acylates lysophospholipids with PUFAs, and thus an essential component of ferroptosis [13]. Indeed, LPCAT3 KD rescued the increase in 4-HNE induced by GPx4 KD (Fig. 4A and Supplemental Fig. S5A). Remarkably, deletion of LPCAT3 KD completely restored the reduction in myotube diameter induced by GPx4 KD (Fig. 4D&E). Similarly, LPCAT3 deletion also prevented LOOH and cell death induced by erastin (Fig. 5F-H and Supplemental Fig. S5B&C). These findings indicate that muscle atrophy induced by loss of GPx4 or erastin treatment is due to the accumulation of LOOH and not other peroxides.

**Fig. 4.**
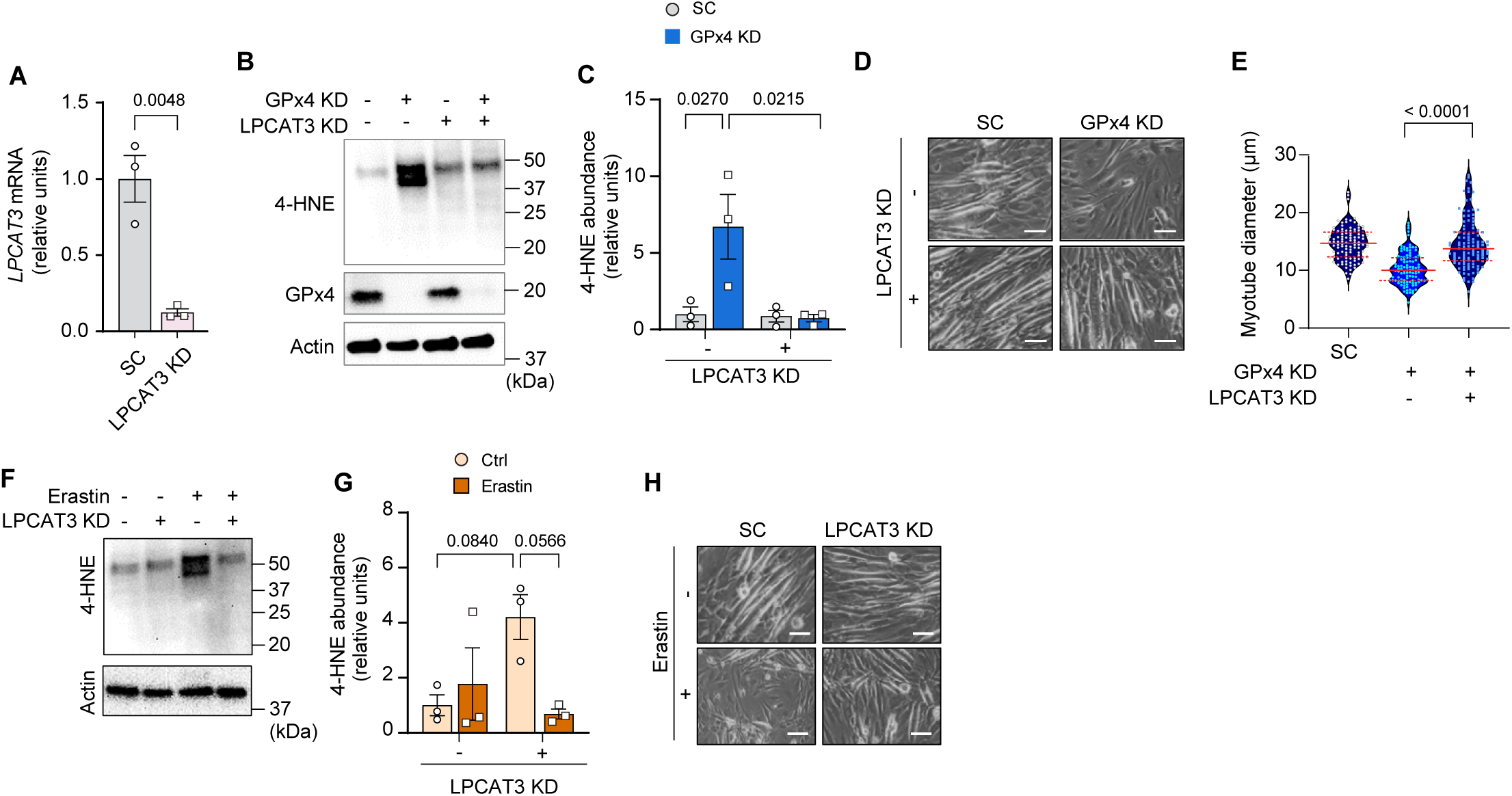
Suppression of PUFA incorporation prevents LOOH-induced myotube atrophy. **(A)** mRNA levels of LPCAT3 in C2C12 myotubes with or without LPCAT3 KD (*n =* 3 per group). **(B)** Immunoblotting of 4-HNE, GPx4 and Actin protein in C2C12 myotubes with or without GPx4 KD and/or LPCAT3 KD. **(C)** Quantification of 4-HNE proteins in C2C12 myotubes with or without LPCAT3 KD and/or GPx4 KD (*n* = 3 per group). **(D,E)** Representative images **(D)**, and quantification of myotube diameter **(E)** from C2C12 myotubes with or without GPx4 KD and/or without LPCAT3 KD (*n =* 104-114 per group). Scale bar, 100 μm. **(F,G)** Immunoblotting **(F)** and quantification **(G)** of 4-HNE from C2C12 myotubes with or without LPCAT 3 KD and/or erastin (*n =* 3 per group). **(H)** Representative images from C2C12 myotubes with or without LPCAT3 KD and/or erastin (*n =* 3 independent repeats). Scale bar, 100 μm. Data are shown as the mean ± SEM. Statistical analyses in **(A)** were performed with an unpaired two-tailed t-test. Statistical analysis in **(C)** and **(G)** were performed with a two-way ANOVA and multiple comparisons were performed using Tukey’s multiple comparisons tests. Statistical analyses in **(E)** was performed with a one-way ANOVA with Dunnett’s multiple comparisons test.

**Fig. 5.**
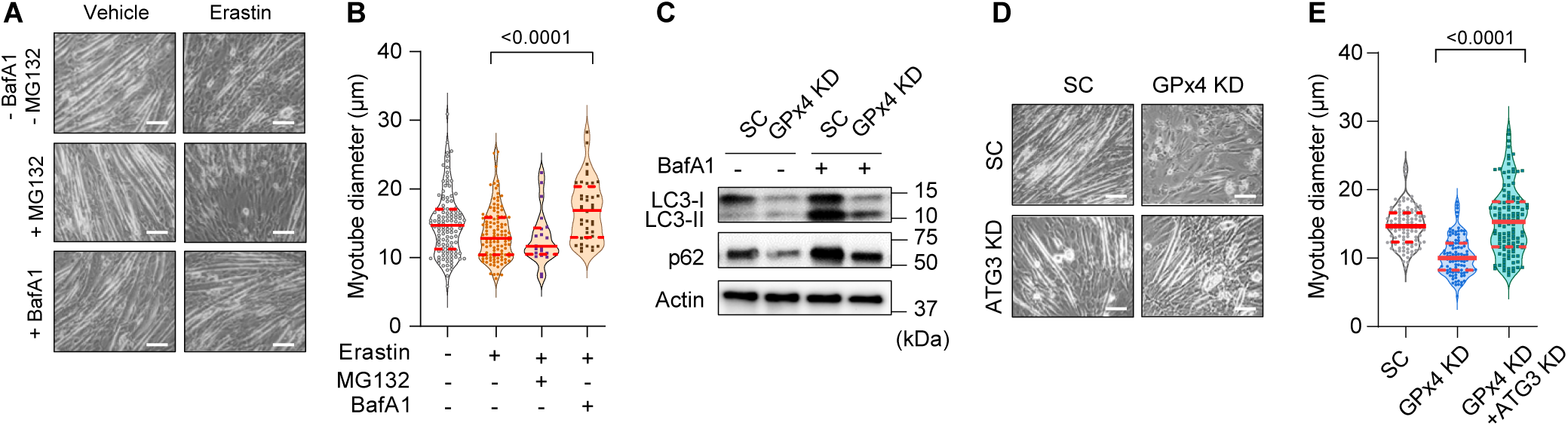
Suppression of autophagy-lysosome axis prevents LOOH-induced myotube atrophy. **(A,B)** Representative images **(A)**, and quantification of myotube diameter **(B)** from erastin-stimulated C2C12 myotubes with proteasomal inhibitor MG132 or lysosomal inhibitor BaFA1. (*n =* 21-120 per group). Scale bar, 100 µm. **(C)** Autophagic flux analyses immunoblotting for LC3-I, LC3-II, p62, and actin in SC or GPx4 KD C2C12 myotubes with or without BafA1. **(D,E)** Representative images **(D)** and quantification of myotube diameter **(E)** from C2C12 myotubes with or without GPx4 KD and/or ATG3 KD) (*n =* 104-121 per group). Data are shown as the mean ± SEM. Statistical analyses in **(B,E)** were performed with a one-way ANOVA with Dunnett’s multiple comparisons test.

What is the mechanism by which LOOH promotes muscle atrophy? C2C12 myotubes were pretreated with Bafilomycin A1 (BafA1) or MG132 prior to erastin incubation to determine whether LOOH increases protein degradation in a lysosomal- or proteasomal-dependent manner, respectively. Erastin-induced reduction in myotube diameter was suppressed with BafA1, but not with MG132 (Fig. 5A&B and Supplemental Fig. S6A), suggesting that the lysosome mediates protein degradation by LOOH [23]. We also reproduced these findings with RSL3 treatment (Supplemental Fig. S6B&C). How does LOOH, a lipid molecule, promote lysosomal-degradation? Upstream of the lysosome, autophagosome formation is mediated by a lipidation of LC3 by ATG3 [24]. Thus, we hypothesized that LOOH may affect the lipidation of LC3. Indeed, GPx4 KD drastically reduced the protein content of p62, LC3-I and LC3-II (Fig. 5C and Supplemental Fig. S6D-H), potentially suggesting that LOOH may accelerate lysosomal degradation by affecting LC3 lipidation. To test this possibility, we performed a targeted deletion of ATG3 in vitro. Indeed, ATG3 KD completely rescued the reduction in myotube diameter induced by GPx4 KD (Fig. 5D&E and Supplemental Fig. S6I-L).

Leveraging these findings, we generated mice with skeletal muscle-specific tamoxifen-inducible ATG3 knockout (ATG3-MKO) (Fig. 6A&B and Supplemental Fig. S7A&B) and studied them with or without HU (Supplemental Fig. S7C-E). Loss of muscle ATG3 was protective from disuse-induced atrophy (Fig. 6C and Supplemental Fig. S7F) and weakness (Fig. 6D and Supplemental Fig. S7G) which can be explained by greater myofiber cross-sectional area (Fig. 6E&D and Supplemental Fig. S7I). Thus, suppression of autophagy is sufficient to attenuate disuse-induced muscle atrophy and weakness.

**Fig. 6.**
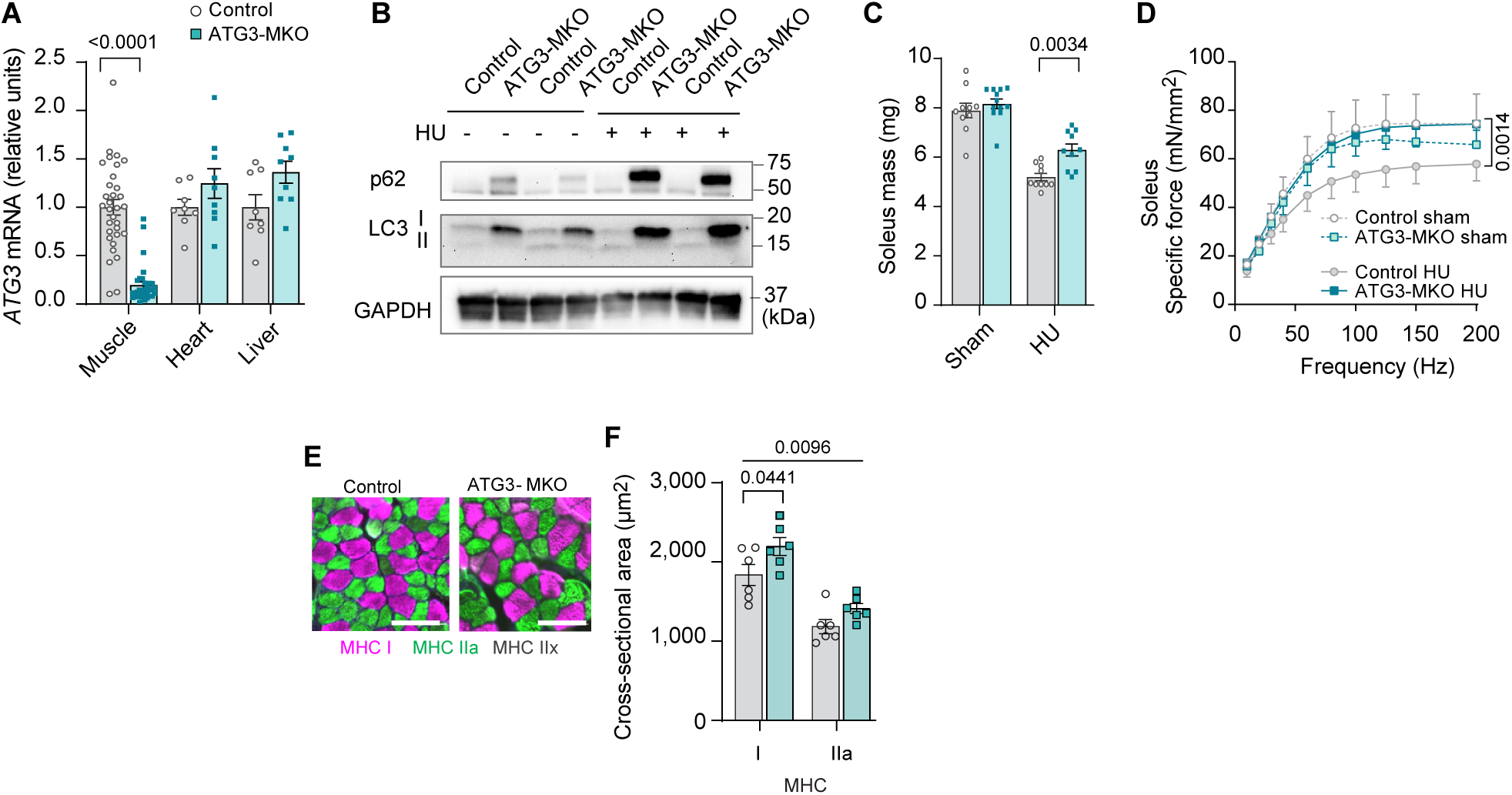
Muscle-specific ATG3 deletion attenuates disuse-induced atrophy and weakness. **(A)** mRNA levels of ATG3 (*n =* 8-32 per group) from young control or ATG3-MKO mice. (**B)** Immunoblotting of p62, LC3, and GAPDH proteins from skeletal muscles from control and ATG3-MKO mice. **(C)** Soleus muscle mass from control or ATG3-MKO mice (*n =* 10-12 per group). **(D)** Force-frequency curve from young control or ATG3-MKO mice (*n =* 9-12 per group). **(E,F)** Representative images of MHC immunofluorescence **(E)** and muscle fiber CSA by fiber type **(F)** of soleus muscles in young control or ATG3-MKO mice with HU (*n =* 6 per group). Data are shown as the mean ± SEM. Statistical analyses in **(A)** were performed with an unpaired two-tailed t-test. Statistical analyses in **(C,D)** and **(F)** were performed with a two-way ANOVA and multiple comparisons were performed using Tukey’s multiple comparisons tests.

We initially hypothesized that lysosomal-degradation mediates LOOH-induced protein degradation to contribute to muscle atrophy. However, further assessment of muscle LOOH illuminated a more complex interaction between the lysosome and LOOH [25, 26]. Unexpectedly, quantification of 4-HNE revealed that inhibition of the autophagy-lysosome axis by ATG3 deletion or BafA1 was sufficient to inhibit LOOH induced by GPx4 deletion, erastin, or RSL3 (Fig. 7A-D and Supplemental Fig. S8A-D). These findings suggest that the autophagy-lysosome axis is essential for LOOH amplification, in addition to its potential role in mediating protein degradation downstream. Indeed, immunofluorescence experiments revealed that 4-HNE is highly co-localized to LAMP2 (Fig. 7E and Supplemental Fig. S8E), consistent with the notion that the lysosome is necessary for LOOH propagation. To support this idea, an increase in LOOH by hydrogen peroxide or carmustine (agents that increase pan oxidative stress without acting on GPx4 directly) was completely inhibited by lysosomal inhibition (Fig. 7F-H and Supplemental Fig. S8F&G). Together, these observations suggest that the propagation of LOOH may be mediated by the lysosome (Supplemental Fig. S8H).

**Fig. 7.**
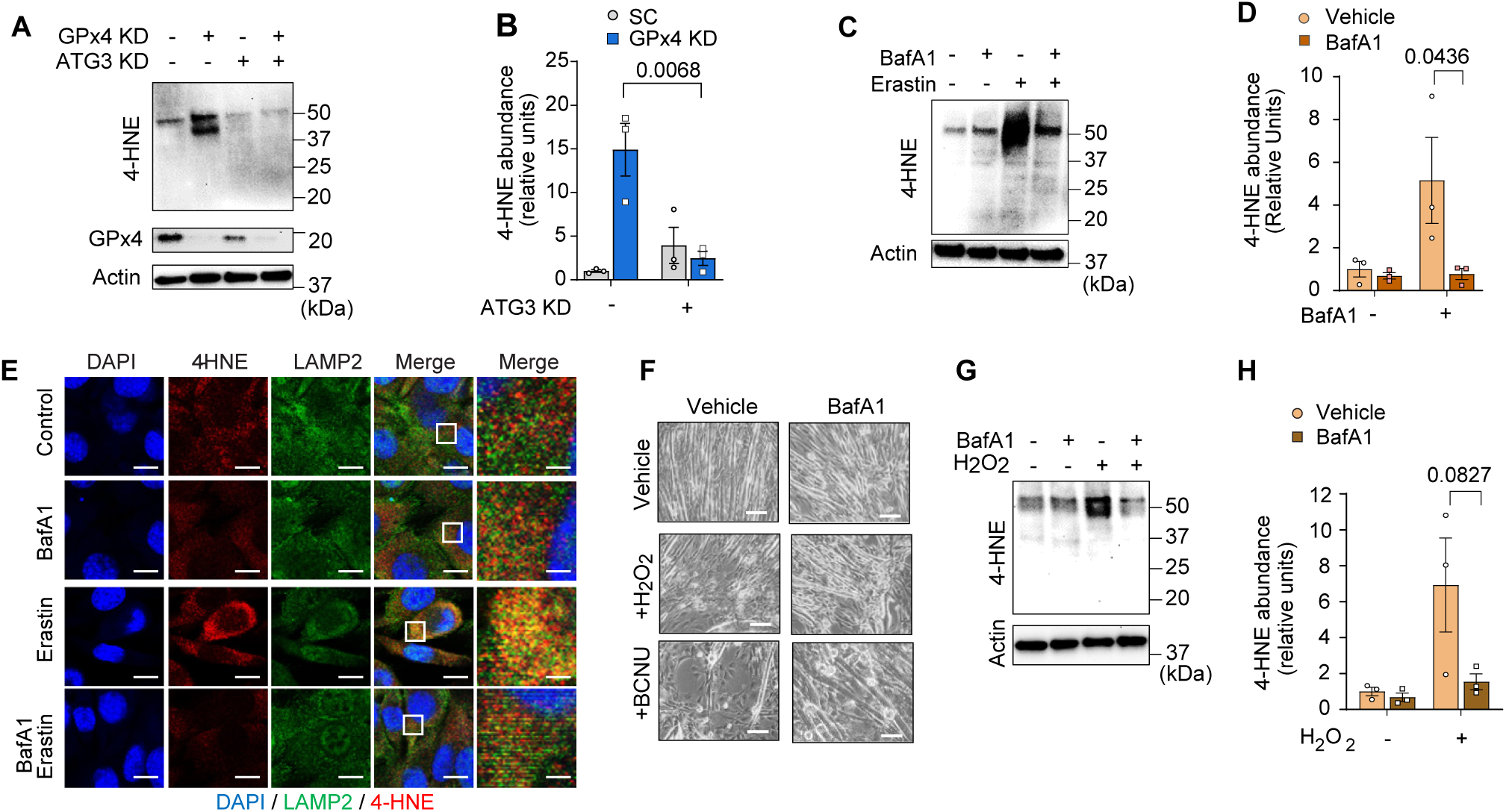
Inhibition of autophagy-lysosome axis prevents accumulation of LOOH. **(A,B)** Immunoblotting **(A)** and quantification **(B)** of 4-HNE, GPx4, and actin in C2C12 myotubes with or without GPx4 KD and/or ATG3 KD. **(C,D)** Immunoblotting **(C)** and quantification **(D)** of 4-HNE protein from C2C12 myotubes with or without erastin and/or BaFA1 (*n =* 3 per group). **(E)** Confocal fluorescence microscope images of erastin-stimulated myotubes with or without BaFA1. Scale bar, 10µm. Boxed regions are shown enlarged at far right. Scale bar, 2.5 µm. **(F,G,H)** Representative images (**F**), immunoblotting of 4-HNE and actin (**G**), and quantification (**H**) of H_2_O_2_ or BCNU-stimulated C2C12 myotubes with or without BaFA1. Scale bar, 100 µm. Data are shown as the mean ± SEM. Statistical analyses in **(B,D)** and **(H)** were performed with a two-way ANOVA and Tukey’s multiple comparison test.

Inhibition of autophagy suppressed lysosomal-degradation and LOOH to attenuate muscle atrophy. We next tested whether suppression of LOOH would be sufficient to ameliorate skeletal muscle atrophy. We studied young (4 months) and old (20 months) global GPx4-overexpressing (GPx4Tg) mice [27] with or without HU (Supplemental Fig. S9A-E). Strikingly, both young and old GPx4Tg mice were resistant to disuse-induced muscle atrophy (Fig. 8A and Supplemental Fig. 9F&G). Perhaps even more impactful was the effect of GPx4 overexpression on skeletal muscle force-generating capacity such that, in both young and old, GPx4 overexpression robustly protected mice from muscle weakness induced by HU (Fig. 8B and Supplemental Fig. S9H-J). These findings are in contrast to our experiments in GPx4^+/−^ mice where muscle mass phenotype was only present in the young mice (Fig. 3A) and no phenotype on muscle strength (Fig. 3C). Consistent with the notion that GPx4 overexpression acts on LOOH, HU-induced increase in 4-HNE was completely suppressed in GPx4Tg mice (Fig. 8C and Supplemental Fig. S9K). We also found that the protection from muscle atrophy was explained by greater myofiber cross-sectional area regardless of fiber-type (Fig. 8D&E and Supplemental Fig. S9L&M).

**Fig. 8.**
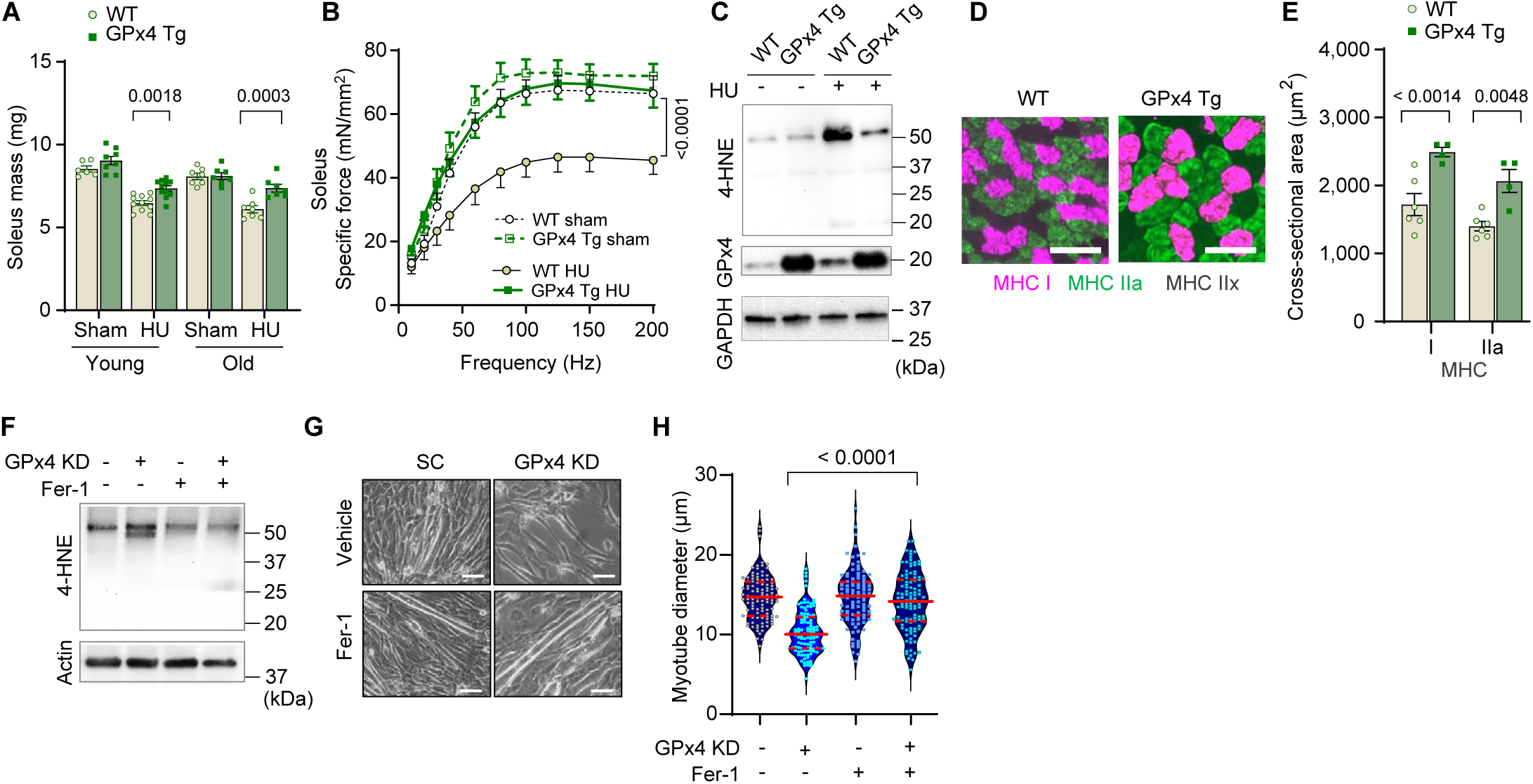
Overexpression of GPx4 ameliorates disuse-induced muscle atrophy and weakness in young and old mice. **(A)** Soleus muscle mass from young or old WT or GPx4Tg mice with or without HU (*n =* 6-11 per young group, *n =* 7 per old group). **(B)** Force-frequency curve from old WT or GPx4Tg mice (*n =* 5-7 per group). **(C)** Immunoblotting of muscle 4-HNE from old WT or GPx4Tg mice. **(D,E)** Representative images of MHC immunofluorescence **(D)** and muscle fiber CSA by fiber type **(E)** for soleus muscles in old WT or GPx4Tg mice with HU (*n =* 4-6 per group). **(F)** Immunoblotting of 4-HNE, and actin in C2C12 myotubes with or without GPx4 KD and/or Ferrostatin-1. **(G,H)** Representative images **(G)**, and quantification of myotube diameter **(H)** from C2C12 myotubes with GPx4 KD and/or Ferrostatin-1 treatments (*n =* 102-110 per group). Scale bar, 100 μm. Data are shown as the mean ± SEM. Statistical analyses in **(H)** was performed with a one-way ANOVA with Dunnett’s multiple comparisons test, and statistical analyses in **(A,B)** and **(E)** were performed with a two-way ANOVA and were performed using Tukey’s **(B,E)** or Sidak’s **(A)** multiple comparisons tests.

Next, we explored opportunities to pharmacologically suppress LOOH to prevent muscle atrophy. Ferrostatin-1 inhibits the propagation of lipid peroxidation and is widely used to study LOOH [28, 29]. Indeed, incubation of cells with ferrostatin-1 was sufficient to suppress LOOH induced by GPx4 KD (Fig. 8F and Supplemental Fig. S10A) concomitant with protection from myotube atrophy (Fig. 8G&H and Supplemental Fig. S10B&C). Nevertheless, ferrostatin-1 is currently not an FDA approved drug with uncertainty surrounding safety. Thus, we tested L-carnosine, a dipeptide composed of beta-alanine and L-histidine that has the ability to scavenge reactive lipid aldehydes formed from LOOH [30, 31]. Rather than acting to suppress the lipid peroxidation process, L-carnosine binds to reactive lipid aldehydes to neutralize carbonyl stress. Similar to ferrostatin-1, L-carnosine was sufficient to suppress 4-HNE and rescue cell death induced by GPx4 KD (Supplemental Fig. S10D-F) or erastin (Supplemental Fig. S10G-I). Leveraging these data, we performed a preclinical trial for L-carnosine provided in drinking water ad lib (80 mM) in young wildtype C57BL6/J mice. L-carnosine treatment did not alter body mass, body composition, food intake, and water intake (Supplemental Fig. S11A-D), and successfully suppressed muscle 4-HNE induced by HU (Supplemental Fig. S11E&F). Remarkably, mice provided with L-carnosine were partly protected from disuse-induced muscle atrophy (Supplemental Fig. S11G).

In humans, L-carnosine is rapidly degraded by a circulating carnosinase [32] that may render oral carnosine treatment ineffective. In contrast, N-acetylcarnosine has a longer half-life and may be a more effective reagent in humans thus improving its translational potential. Similar to ferrostatin-1 and L-carnosine, N-acetylcarnosine also prevented 4-HNE and cell death induced by GPx4 KD (Supplemental Fig. S12A-C) or erastin (Supplemental Fig. S12D-F). Thus, we proceeded with a preclinical trial for N-acetylcarnosine in drinking water (80 mM, Fig. 9A) in young (4 months, C57BL6/J; Jax colony) and old (20 months, C57BL/6; NIA rodent colony) wildtype mice. Similar to L-carnosine treatment, N-acetylcarnosine did not alter body mass, body composition, food intake, or water intake (Supplemental Fig. S13A-E), and successfully suppressed muscle 4-HNE (Fig. 9B and Supplemental Fig. S13F&G). Strikingly, similar to our findings in GPx4Tg mice, N-acetylcarnosine ameliorated muscle atrophy (Fig. 9C and Supplemental Fig. S13H) and weakness (Fig. 9D and Supplemental Fig. S13I-K) in both young and old mice. Protection from muscle atrophy was similarly explained by greater myofiber cross-sectional area regardless of fiber-type (Fig. 9E&F and Supplemental Fig. S13L&M).

**Fig. 9.**
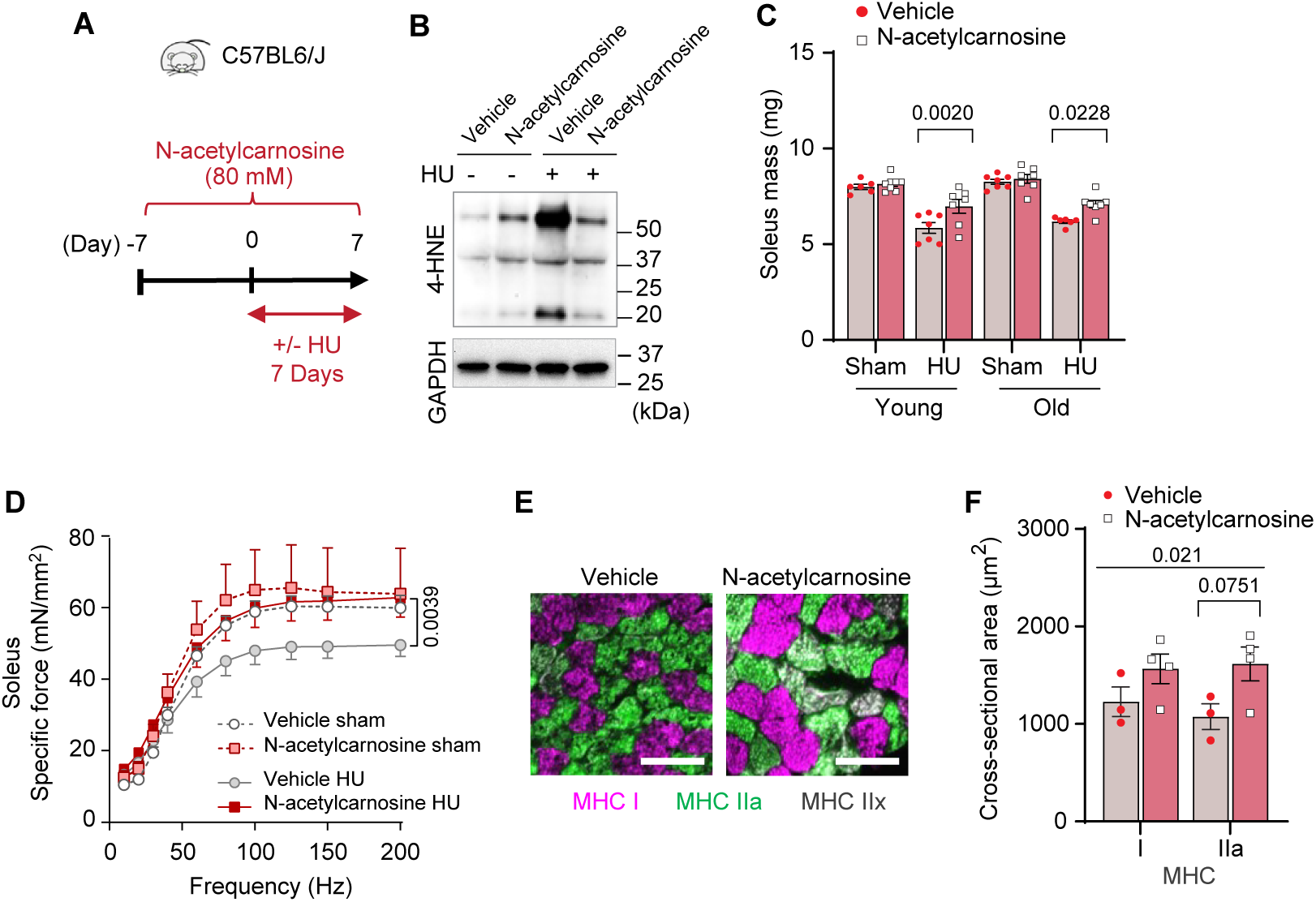
Pharmacologic suppression of carbonyl stress ameliorates muscle atrophy and weakness in young and old mice. **(A)** Schematic illustration of the protocol for administration of N-acetylcarnosine in vivo. **(B)** Immunoblotting of muscle 4-HNE from N-acetylcarnosine treatment in old mice. **(C)** Soleus muscle mass from young or old mice with or without N-acetylcarnosine treatment (*n =* 6-8 per young group, *n =* 7 per old group). **(D)** Force-frequency curve from N-acetylcarnosine study in old mice (*n =* 4-5 per group). **(E,F)** Representative images of MHC immunofluorescence **(E)** and muscle fiber CSA by fiber type **(F)** for soleus muscles (*n =* 3-4 per group) in old mice from the N-acetylcarnosine study. Data are shown as the mean ± SEM. Statistical analyses in **(C,D)** and **(F)** were performed with a two-way ANOVA and were performed using Tukey’s **(C,F)** or Sidak’s **(D)** multiple comparisons tests.

## Discussion

The current findings demonstrate a novel mechanism that indicate LOOH as the key downstream molecule by which oxidative stress promotes muscle atrophy and weakness. Skeletal muscle LOOH was robustly upregulated with aging and disuse, and genetic or pharmacologic neutralization of LOOH and their secondary reactive lipid aldehydes was sufficient to rescue muscle atrophy and weakness. In particular, N-acetylcarnosine treatment shows a potent effect in preserving muscle mass and strength with disuse in both young and old mice, informing the potential trial to utilize this compound to ameliorate loss of muscle function in humans.

During the preparation of this manuscript, Van Remmen and colleagues published a complementary study demonstrating that liproxstatin-1 can suppress denervation-induced skeletal muscle atrophy [33]. Denervation and HU elicits different but overlapping response in myofibers, and our studies demonstrate that their effects to drive skeletal muscle atrophy might converge on lipid peroxidation. Like ferrostatin-1, liproxstatin-1 acts to suppress the propagation of lipid peroxidation rather than acting directly on LOOH. Nevertheless, in vivo liproxstatin-1 treatment was highly effective in suppressing denervation-induced LOOH as well as reactive lipid aldehydes 4-HNE, suggesting that targeting lipid peroxidation is likely an equally effective strategy to suppress LOOH production in skeletal muscle. Conversely, our data with GPx4 overexpression and N-acetylcarnosine treatment indicate that the effect of lipid peroxidation to promote muscle atrophy is mediated by LOOH and their lipid reactive aldehydes. Neither liproxstatin-1 nor ferrostatin-1 are currently FDA approved, but it is worthwhile to consider these drugs along with N-acetylcarnosine as potential therapeutics to treat muscle atrophy.

Sarcopenia is an age-associated decline in muscle mass and strength, that occurs due to a combination of inactivity, injury, and/or surgery, in addition to the biological consequences of aging itself. In the current study, mice were studied at 4 or 20 months of age. While not statistically compared directly (these experiments were not performed side-by-side), skeletal muscle mass at 20 months of age was not significantly lower compared to those at 4 months of age. Thus, the current data is unclear whether targeting LOOH prevents the loss of muscle mass due to the biological effect of aging in the absence of HU. We chose to study mice with 20 months of age for two reasons. First, mice greater than 20 months of age do not tolerate the HU intervention well, often resulting in their inability to consume food or water. Because disuse is an integral component of human aging, we wanted to study how muscles from old mice respond to disuse. This therefore compromised our ability to study sarcopenia without disuse. Second, while muscle mass was not diminished at 20 months of age, skeletal muscle force-generating capacity was lower in 20 months old mice compared to the 4 months old mice, particularly in the extensor digitorum longus (EDL) muscles. In GPx4Tg mice, age-associated decrease in muscle strength (in non-HU mice) appeared to be rescued, while short-term treatment with N-acetylcarnosine had no effect. We are currently following up on these results with a long-term treatment of N-acetylcarnosine in 24 months old mice to see if such intervention might alleviate the loss of muscle mass and strength associated with age in absence of the HU intervention.

We initially set out to investigate the role of LOOH in age- and disuse-induced skeletal muscle atrophy, while measuring force-generating capacity as a secondary outcome. However, in all experimental models in which accumulation of muscle LOOH was suppressed (young and old GPx4Tg mice, young and old mice with N-acetylcarnosine, and young ATG3-MKO mice), force-generating capacity (i.e., specific force normalized to cross-sectional area) was more robustly rescued compared to skeletal muscle mass. This suggests the role of LOOH to induce muscle weakness independent of muscle atrophy, and likely independent of muscle protein degradation. While out of the scope for the current study, it would be important to determine whether reactive lipid aldehydes induced by aging or disuse preferentially bind to enzymes of skeletal muscle contraction to compromise their activities. As described in the previous paragraph, aging promoted muscle weakness prior to atrophy. Similarly, muscle atrophy induced by cancer cachexia is also preceded by muscle weakness [34]. Thus, these observations highlight the need to better study the mechanisms that regulate force-generating capacity independent of muscle mass.

In conclusion, we provide evidence that LOOH contributes to the loss of muscle mass and strength associated with age and disuse. Neutralization of LOOH, particularly their reactive lipid aldehyde byproducts, attenuates muscle atrophy and weakness. The mechanisms by which LOOH contributes to these phenotypes are not entirely clear, but they include protein degradation mediated by the autophagy-lysosomal axis, as well as loss in the force-generating capacity that is likely mediated by carbonyl stress. Last, but not least, these promising observations inform a potential clinical trial to test the efficacy of N-acetylcarnosine treatment in ameliorating muscle atrophy in humans.

## Methods

### Key Resources

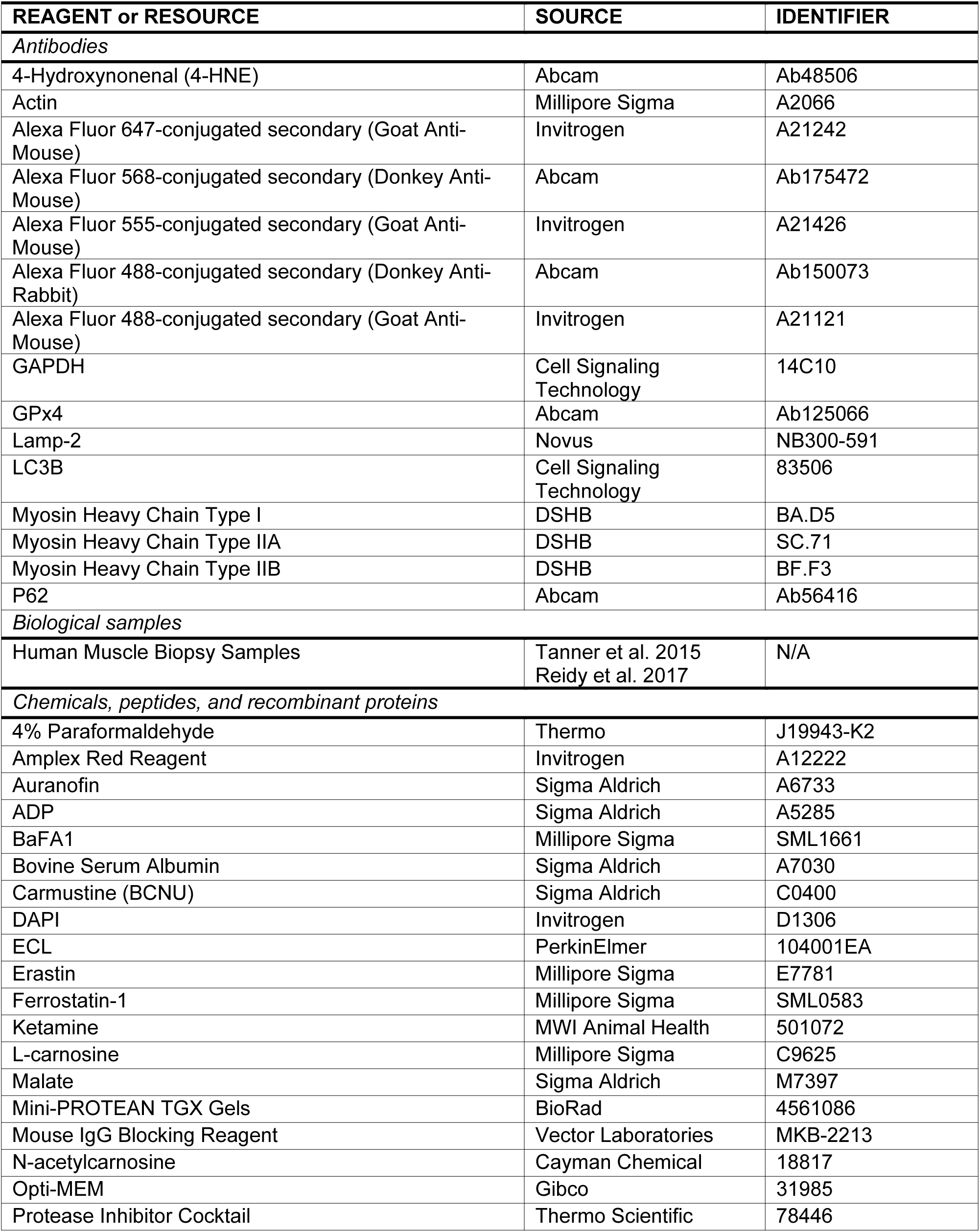
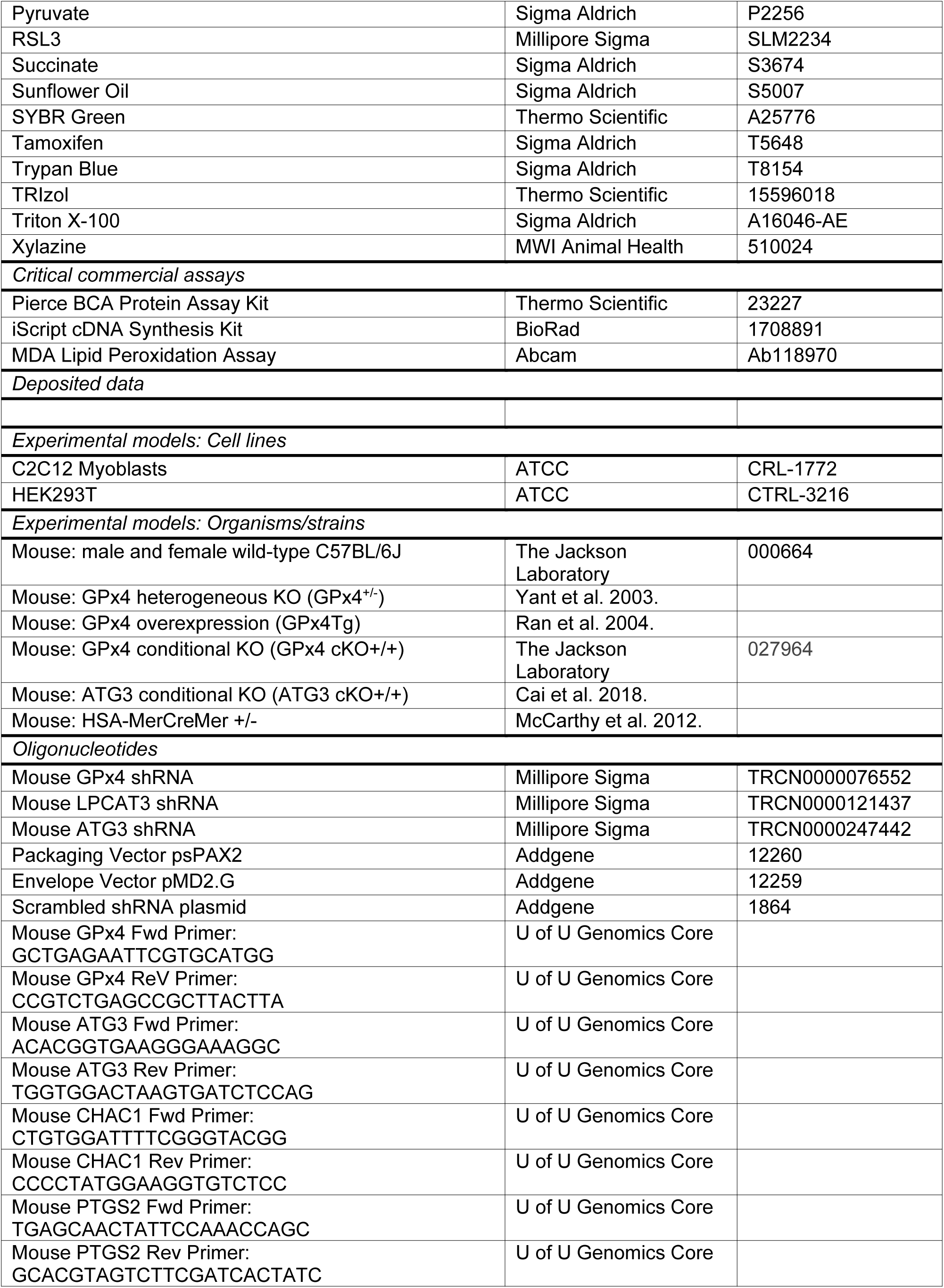
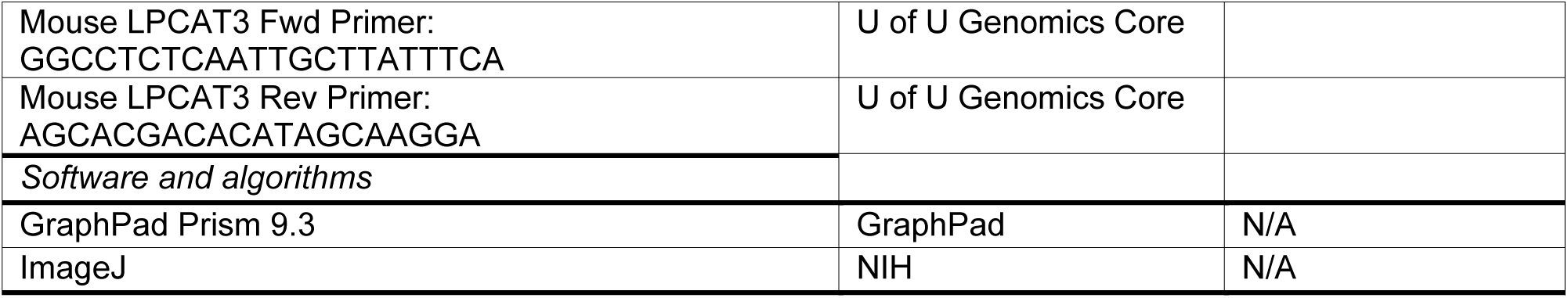

### Animal models

GPx4+/− and GPx4Tg mice were generated previously [16, 27]. Conditional GPx4 knockout (GPx4cKO+/+) mice were acquired from Jackson Laboratory (Stock No: 027964) [18]. Conditional ATG3 knockout (ATG3cKO+/+) mice were previously described [35]. GPx4cKO+/+ mice or ATG3cKO+/+ mice were then crossed with tamoxifen-inducible, skeletal muscle–specific Cre recombinase (HSA-MerCreMer+/−) mice [36] to generate GPx4cKO+/+; HSAMerCreMer−/− (control) and GPx4cKO+/+; HSA-MerCreMer+/− (skeletal muscle–specific GPx4 knockout; GPx4-MKO) mice or ATG3cKO+/+; HSAMerCreMer−/− (control) and ATG3cKO+/+; HSA-MerCreMer+/− (ATG3-MKO) mice. Tamoxifen-injected (7.5 μg/g body mass, 5 consecutive days) littermates were used. Mice were maintained on a 12-hour light/12-hour dark cycle in a temperature-controlled room. Body composition measurements were taken immediately before terminal experiments with a Bruker Minispec MQ20 nuclear magnetic resonance (NMR) analyzer (Bruker, Rheinstetten, Germany). All mice were bred onto C57BL/6J background and were born at normal Mendelian ratios. Body mass were measured every day during HU. All protocols were approved by Institutional Animal Care and Use Committees at the University of Utah.

### Hindlimb unloading

Mice underwent 1, 7, or 14 days of HU (2 mice/cage) using a previously described protocol [14, 15] based on the traditional Morey-Holton design to study disuse atrophy in rodents. Along with daily monitoring of body mass, food intake was monitored every other day to ensure that the mice did not experience excessive weight loss due to malnutrition or dehydration. Following 1, 7, or 14 days of HU, mice were fasted for 4 h and given an intraperitoneal injection of 80 mg/kg ketamine and 10 mg/kg xylazine, after which tissues were harvested. Extensor digitorum longus (EDL), and soleus (SOL) were carefully dissected for weight measurements.

### Muscle force generation

Force-generating properties of soleus and EDL muscles were measured as previously described [37, 38]. Briefly, soleus/EDL muscles were sutured at each tendon, and muscles were suspended at optimal length (Lo), which was determined by pulse stimulation. After Lo was identified, muscles were stimulated (0.35 seconds, pulse width 0.2 milliseconds) at frequencies ranging from 10 to 200 Hz. Muscle length and mass were measured to quantify cross-sectional area for force normalization.

### Quantitative reverse transcription PCR

Samples were homogenized in TRIzol reagent (Life Technologies) to extract total RNA. One microgram RNA was reverse-transcribed using an IScript cDNA synthesis kit (Bio-Rad). Reverse transcription PCR (RT-PCR) was performed with the Viia 7 Real-Time PCR System (Life Technologies) using SYBR Green reagent (Life Technologies). All data were normalized to ribosomal L32 gene expression and were normalized to the mean of the control group. Primers were based on sequences in public databases.

### Western blot

Whole muscle or cells were homogenized, and western blots were performed as previously described [14]. Protein homogenates were analyzed for abundance of phosphorylated 4-hydroxynonenal (4-HNE; ab48506; Abcam), GPx4 (ab125066, Abcam), actin (A2066, MilliporeSigma), GAPDH (14C10, Cell Signaling Technology), p62 (ab56416, Abcam), LC3B (83506, Cell Signaling Technology).

### Mass spectrometry

Oxidolipidomics samples were analyzed on the SCIEX 7500 system coupled with ExionLC (SCIEX, Concord, Canada) using multiple reaction monitoring (MRM) analysis. Mobile phase A is composed of 93:7 acetonitrile:dichloromethane containing 2mM ammonium acetate and mobile phase B is composed of 50:50 acetonitrile:water containing 2mM ammonium acetate. A Phenomenex Luna® NH2 3 µm particle size (4.6×150mm) was used for separation and column temperature was kept at 40°C. The total flow rate is 0.7 mL/min with a total run time of 17-minutes. Samples were extracted using the Bligh & Dyer method. Lower layer was collected, dried down and resuspended in mobile phase A.

### Cell culture

C2C12 myoblasts were grown and maintained in high-glucose Dulbecco’s modified Eagle’s medium (DMEM), with 10% fetal bovine serum (FBS), and 0.1 % penicillin/streptomycin. Once 90 to 100% confluent, C2C12 cells were differentiated into myotubes with low-glucose DMEM, with l-glutamine and 110 mg/L sodium pyruvate; supplemented with 2% horse serum, and 0.1% penicillin-streptomycin. For experiments with erastin (E7781, MilliporeSigma), Ferrostatin-1 (SML0583, MilliporeSigma), and RSL3 (SML2234, MilliporeSigma), C2C12 myotubes were incubated with either 10 μM erastin/10 µM Ferrostatin-1/5 μM RSL3/ or equal-volume DMSO directly dissolved into medium. For experiments with L-carnosine (C9625, MilliporeSigma), and N-acetylcarnosine (18817, Cayman), C2C12 myotubes were incubated with 10 mM of L-carnosine/N-acetylcarnosine directly dissolved into medium.

### Lentivirus-mediated knockdown of GPx4/LPCAT3/ATG3

Lentivirus-mediated knockdown of experiments were performed as previously described [15, 21, 39]. Vectors were decreased using pLKO.1 lentiviral-RNAi system. Plasmids encoding short hairpin RNA (shRNA) for mouse GPx4 (shGPx4: TRCN0000076552), mouse LPCAT3 (shLPCAT3: TRCN0000121437), and mouse ATG3 (shATG3: TRCN0000247442) were obtained from MilliporeSigma. Packaging vector psPAX2 (ID 12260), envelope vector pMD2.G (ID 12259), and scrambled shRNA plasmid (SC: ID 1864) were obtained from Addgene. HEK293T cells in 10 cm dishes were transfected using 50 μL 0.1% polyethylenimine, 200 μL 0.15 M sodium chloride, and 500 μL Opti-MEM (with HEPES, 2.4 g/L sodium bicarbonate, and l-glutamine; Gibco 31985) with 2.66 μg of psPAX2, 0.75 μg of pMD2.G, and 3 μg of either scrambled or GPx4/LPCAT3/ATG3 shRNA plasmids. After 48 hours, growth medium was collected, filtered using 0.22 μm vacuum filters, and used to treat undifferentiated C2C12 cells for 48 hours. To ensure that only cells infected with shRNA vectors were viable, cells were selected with puromycin throughout differentiation.

### Measurements of myotube diameter

Images of myotubes were visualized at ×20 magnification using an inverted light microscope and captured with a camera (DP74, Olympus). Myotube diameter was measured for at least 100 myotubes from 5 random fields in each group using ImageJ software. The average diameter per myotube was calculated as the mean of ten short-axis measurements taken along the length of the myotube.

### Assessment of cell death

Cell death levels were examined by counting the numbers of cells with trypan blue staining. The cells were trypsinized and stained with 0.2% trypan blue for 5 min. Stained and non-stained cells were counted under a microscope using a hemocytometer.

### Immunofluorescence

C2C12 myotubes were fixed with 4% paraformaldehyde for 10 min and permeabilized with 0.2% Triton X-100 for 15 min. After blocking with bovine serum albumin, immunocytochemistry was performed with anti-HNE (ab48506, Abcam), anti-lysosome associated membrane protein 2 (Lamp-2) (NB300-591, Novus), and Alexa Fluor-conjugated secondary antibodies Alexa Fluor® 568 (ab175472, abcam), Alexa Fluor® 488 (ab150073, abcam), and DAPI (D1306, Invitrogen). Images were captured using a 63× 1.4 NA oil immersion objective on a Leica SP5 confocal system (Leica). For an experiment, C2C12 myotubes incubated with erastin with or without pretreatment of BaFA1 (SML1661, MilliporeSigma). Soleus muscles were embedded in optimal cutting temperature (OCT) gel and sectioned at 10 µm with a cryostat (Microtome Plus). The sections underwent blocking for 1 hr with M.O.M. mouse IgG Blocking Reagent (Vector Laboratories, MKB-2213), 1 hr with primary antibodies (BA.D5, SC.71, BF.F3 all at 1:100 from DSHB). Sections were then probed with the following secondary antibodies: Alexa Fluor 647 (1:250; Invitrogen, A21242), Alexa Fluor 488 (1:500; Invitrogen, A21121), and Alexa Fluor 555 (1:500; Invitrogen, A21426). Negative stained fibers were considered to be IIx. Slides were imaged with an automated wide-field light microscope (Nikon Corp.) using a 10x objective lens. Cross-sectional area and fiber type composition was then quantified utilizing ImageJ software.

### Mitochondrial respiration measurements

Mitochondrial O_2_ utilization was measured using the Oroboros O_2_K Oxygraphs, as previously described [14, 15]. Isolated mitochondria were added to the oxygraph chambers containing buffer Z. Respiration was measured in response to the following substrate concentrations: 0.5 mM malate, 5 mM pyruvate, 2 mM ADP, 10 mM succinate, and 1.5 μM FCCP.

### Mitochondrial H_2_O_2_ measurements

Mitochondrial H_2_O_2_ production was measured using the Horiba Fluoromax-4, as previously described [14, 15]. Briefly, skeletal muscle was minced in mitochondria isolation medium (300 mM sucrose, 10 mM HEPES, 1 mM EGTA) and subsequently homogenized using a Teflon glass system. Homogenates were then centrifuged at 800 g for 10 min, after which the supernatant was taken and centrifuged at 12,000 g for 10 min. The resulting pellet was carefully resuspended in mitochondria isolation medium. *J*H_2_O_2_ was measured in buffer Z (MES potassium salt; 105 mM, KCl 30 mM, KH_2_PO_4_ 10 mM, MgCl_2_ 5 mM, and BSA 0.5 mg/ml) supplemented with 10 μM Amplex UltraRed (Invitrogen) and 20 U/mL CuZnSOD in the presence of the following substrates: 10 mM succinate, 100 μM 1,3-bis(2-chloroethyl)-1-nitrosourea (BCNU/carmustine), and 1 μM auranofin. The appearance of the fluorescent product was measured with excitation/emission at 565/600 nm.

### Administration of L-carnosine/N-acetylcarnosine in vivo

Carnosine was administered as previously described [31]. Briefly, young (4-month-old) or old (20-month-old) C57BL/6J mice were supplemented with 80 mM carnosine dissolved in drinking water (pH 7.5) for 2 weeks (1 week of pretreatment and 1 week during HU). Bottles were refreshed two times a week (L-carnosine, C9625, MilliporeSigma), and or everyday (N-acetyl carnosine, 18817, Cayman).

### MDA quantification

MDA content was quantified in fresh gastrocnemius muscles using a lipid peroxidation assay kit (ab118970, Abcam) according to the manufacturer’s instruction. Rates of appearance of MDA-thiobarbituric acid adduct were quantified colorimetrically at 532 nm using a spectrophotometer.

### Statistical analyses

Data are presented as means ± s.e.m. Statistical analyses were performed using GraphPad Prism 7.03. Independent sample t-tests (two-sided) were used to compare two groups. For multiple comparisons, one- or two-way analysis of variance (ANOVA) were performed followed by appropriate post-hoc tests corrected for multiple comparisons. For all tests P < 0.05 was considered statistically significant.

## Acknowledgements

This research is supported by NIH grants DK107397, DK127979, GM144613, AG074535, AG063077, (to K.F.), AG050781 (to M.J.D.), HL122863, AG057006 (E.J.A), AG064078 (to Q.R.), HL149870 (S.B.), HL139451 (to Z.S.M), DK130555 (to A.D.P.), AG073493 (to J.J.P.), American Heart Association grants 915674 (to P.S.), 18PRE33960491 (to A.R.P.V.), and 19PRE34380991 (to J.M.J.), Larry H. & Gail Miller Family Foundation (to P.J.F.), University of Utah Center on Aging Pilot Grant (to K.F.), and Uehara Memorial Foundation (to H.E.). We would like to thank Diana Lim from the University of Utah Molecular Medicine Program for assistance with figures.

## Author contributions

H.E. and K.F. contributed to study concept design and wrote the manuscript. P.S. and J.L.S. contributed to study design and data analysis. M.J.P., J.A.M., and J.E.C. performed mass spectrometry analyses. M.J.D. and Z.S.M. performed human muscle biopsies. H.E., P.S., J.L.S., A.W.L., J.M.J., J.J.P., A.S., and E.R.M. performed mouse experiments. H.E. performed all biochemical assays, body composition measurements and analysis of muscle force production. A.R.P.V. and P.J.F. assisted in muscle functional measurements. P.J.F. assisted in cell culture experiments. E.J.A assisted in L-carnosine and N-acetylcarnosine experiments. S.B. and Q.R. designed and generated mouse models. All contributed to editing the manuscript.

## Materials & Correspondence

Correspondence and material requests should be addressed to K. Funai.

## Supplemental Figures

**Supplemental Fig. S1.**
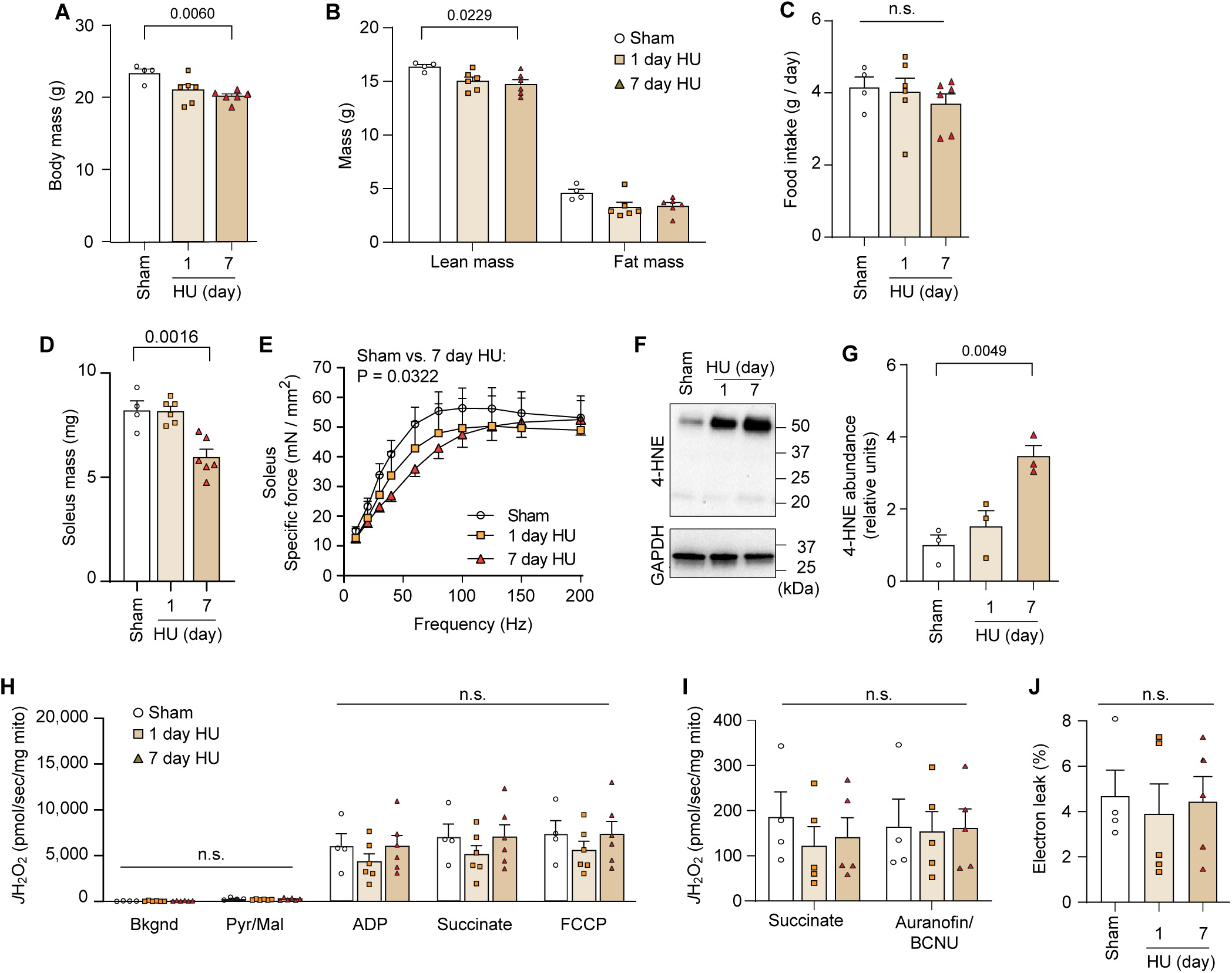
Disuse promotes LOOH in skeletal muscle. **(A,B,C)** Body mass **(A)**, body composition **(B)**, food intake **(C)**, following 1, and 7-days of HU from young WT C57BL/6J mice (*n =* 4 for sham, *n =* 6 for 1-day HU, *n =* 6 for 7-days HU). **(D,E)** Soleus muscle mass **(D)**, force-frequency curve **(E)** in soleus muscle following 1- and 7-days of HU from young WT C57BL/6J mice (*n =* 4 for sham, *n =* 6 for 1-day HU, *n =* 6 for 7-days HU). **(F,G)** Immunoblotting **(F)**, and quantification **(G)** of 4-HNE following 1-day and 7-days of HU (*n =* 3 per group). **(H)** Rates of oxygen consumption measured in isolated skeletal muscle mitochondria with Krebs cycle substrates (*n =* 4 for sham, *n =* 5 for 1-day HU, *n =* 5 for 7-days HU). ADP, adenosine diphosphate; FCCP, carbonyl cyanide-p-trifluoromethoxyphenylhydrazone; Mal, malate; Pyr, pyruvate. **(I)** Rate of mitochondrial H_2_O_2_ production in response to succinate or Aurorafin/BCNU (*n =* 4 for sham, *n =* 5 for 1-day HU, *n =* 5 for 7-day HU). BCNU, 1,3- bis[2-chloroethyl]-1-nitrosourea. **(J)** Percentage of electron leak in muscle mitochondria (*n =* 4 for sham, *n =* 5 for 1-day HU, *n =* 5 for 7-days HU). Data are shown as the mean ± SEM. Statistical analyses in **(A,B,C,D,G)** and **(J)** were performed with a one-way ANOVA and Dunnett’s multiple comparisons tests. Statistical analyses in **(E,H)** and **(I)** were performed with a two-way ANOVA and Tukey’s multiple comparison test.

**Supplemental Fig. S2.**
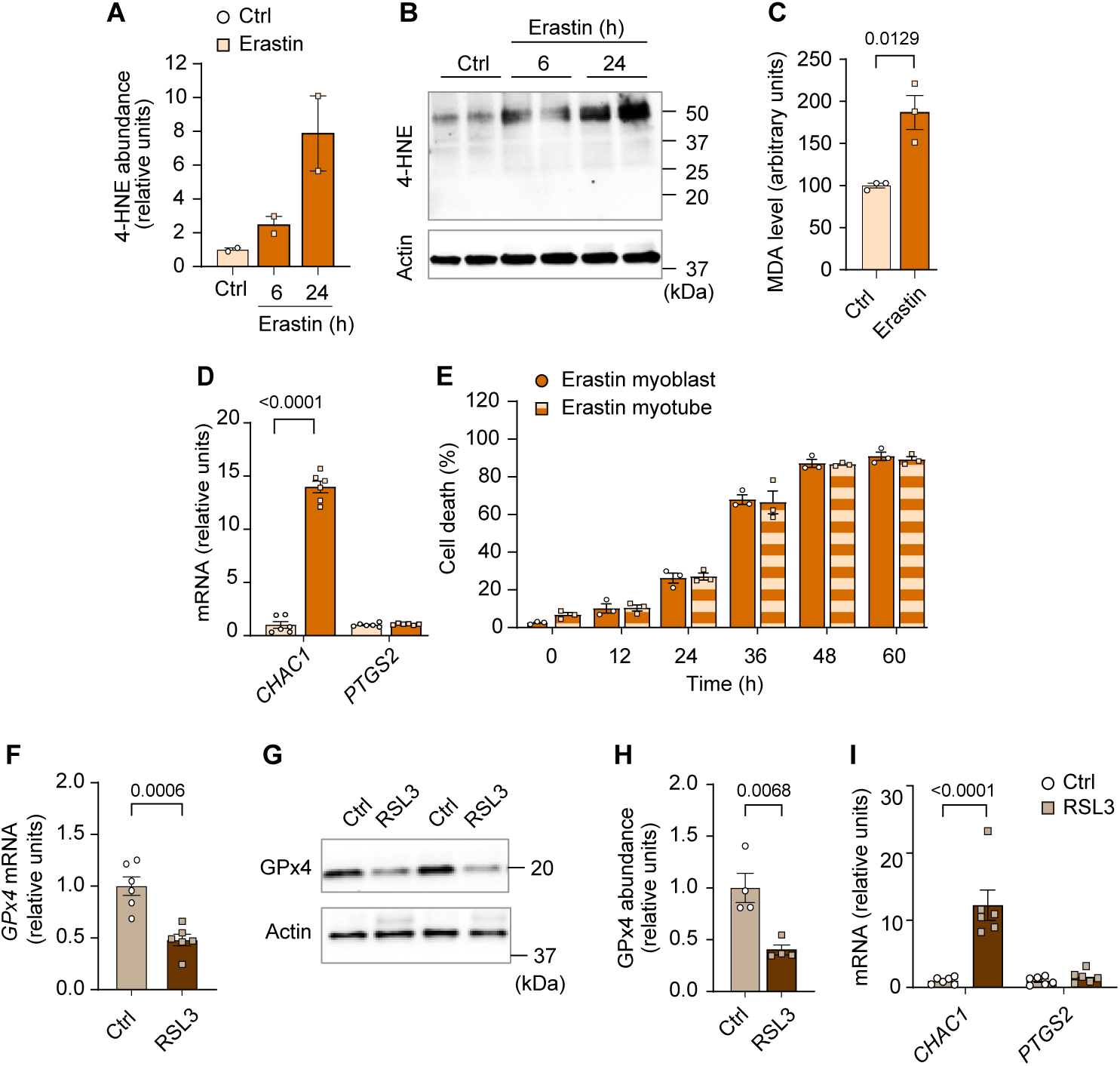
Elevated LOOH is sufficient to promote atrophy in cultured myotubes. **(A,B,C,D,E)** Quantification of 4-HNE **(A)**, immunoblotting of 4-HNE **(B)** proteins, MDA levels **(C)**, mRNA levels for CHAC1 and PTGS2, markers of ferroptosis **(D)**, and cell death levels **(E)** in erastin-stimulated C2C12 myotubes (*n =* 3 independent repeats). **(F,G,H,I)** mRNA levels of GPx4 **(F)**, immunoblotting of GPx4 and actin **(G)**, quantification of GPx4 **(H)**, CHAC1 and PTGS2, markers of ferroptosis (**I**) in RSL3-stimulated C2C12 myotubes (*n =* 6 per group). Data are shown as the mean ± SEM. Statistical analyses in **(C,D,E,F,H,I)** and were performed with an unpaired two-tailed t-test. Statistical analyses in **(A)** were performed with a one-way ANOVA and Dunnett’s multiple comparisons tests.

**Supplemental Fig. S3.**
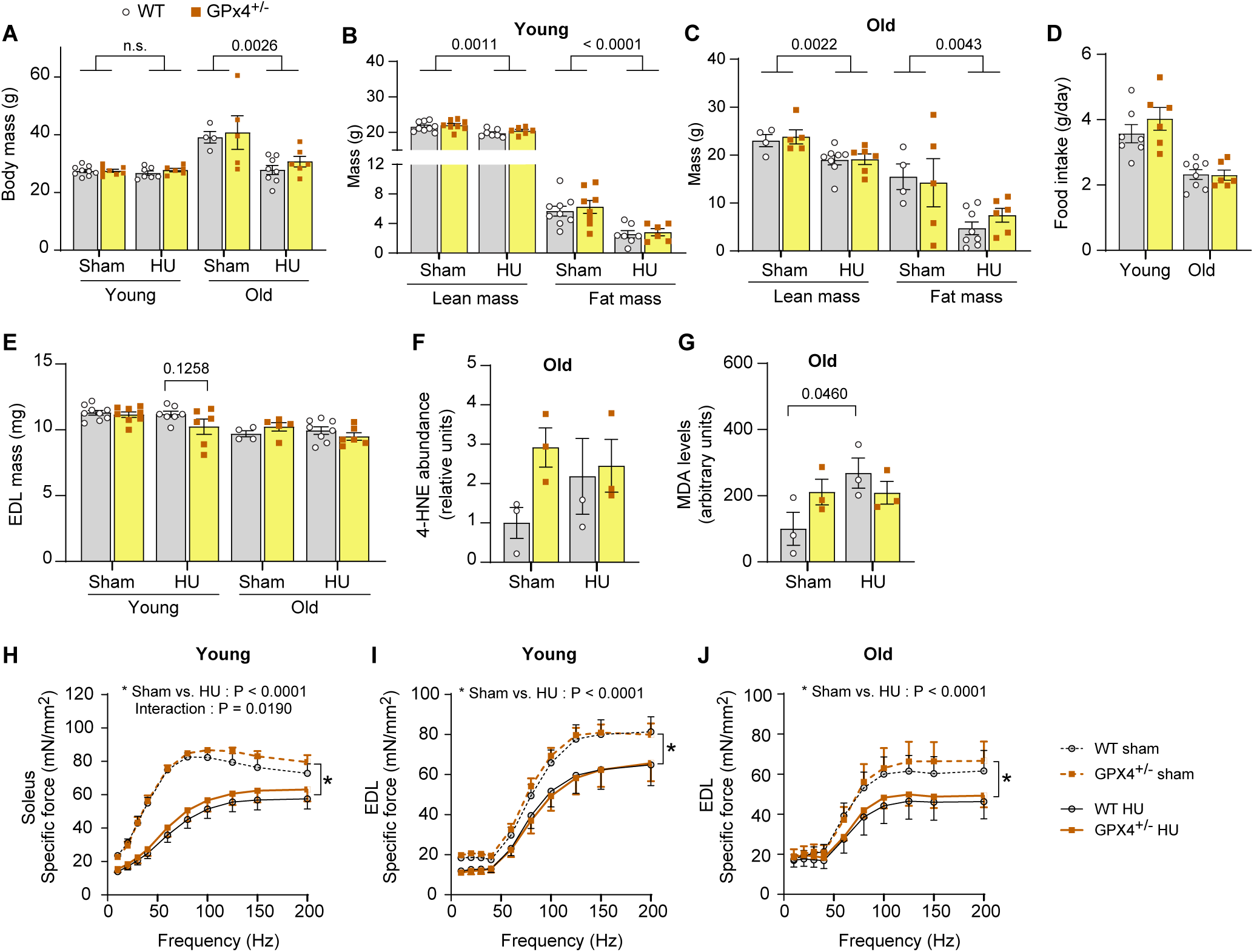
Additional data from GPx4^+/−^ mice. **(A)** Body mass from young and old WT or GPx4^+/−^ mice. **(B,C)** body composition from young (*n =* 9 for WT sham, *n =* 8 for GPx4^+/−^ sham, *n =* 7 for WT HU, *n =* 6 for GPx4^+/−^ HU) **(B)** and old (*n =* 4 for WT sham, *n =* 5 for GPx4^+/−^ sham, *n =* 8 for WT HU, *n =* 6 for GPx4^+/−^ HU) (**C**) WT or GPx4^+/−^ mice. **(D)** Food intake during hindlimb unloading (HU) in young (*n =* 7 for WT, *n =* 6 for GPx4^+/−^) and old (*n =* 8 for WT, *n =* 6 for GPx4^+/−^) WT or GPx4^+/−^ mice. **(E)** EDL muscle mass from young (*n =* 9 for WT sham, *n =* 8 for GPx4^+/−^ sham, *n =* 7 for WT HU, *n =* 6 for GPx4^+/−^ HU) and old (*n =* 9 for WT sham, *n =* 8 for GPx4^+/−^ sham, *n =* 7 for WT HU, *n =* 6 for GPx4^+/−^ HU) WT or GPx4^+/−^ mice. **(F)** Quantification of muscle 4- HNE from old mice (*n =* 3 per group). **(G)** MDA levels from old mice (*n =* 3 per group). (**H**) Force-frequency curve from soleus muscle from young WT or GPx4^+/−^ mice (*n =* 5 for WT sham, *n =* 5 for GPx4^+/−^ sham, *n =* 7 for WT HU, *n =* 5 for GPx4^+/−^ HU). **(I,J)** Force-frequency curve in EDL muscle from young (*n =* 5 for WT sham, *n =* 5 for GPx4^+/−^ sham, *n =* 7 for WT HU, *n =* 5 for GPx4^+/−^ HU) **(I)** and old (*n =* 3 for WT sham, *n =* 4 for GPx4^+/−^ sham, *n =* 6 for WT HU, *n =* 6 for GPx4^+/−^ HU) **(J)** WT or GPx4^+/−^ mice. Data are shown as the mean ± SEM. Statistical analyses in **(A,B,C,D,E,F,G,H,I,J)** were performed with a two-way ANOVA and using Tukey’s multiple comparisons tests.

**Supplemental Fig. S4.**
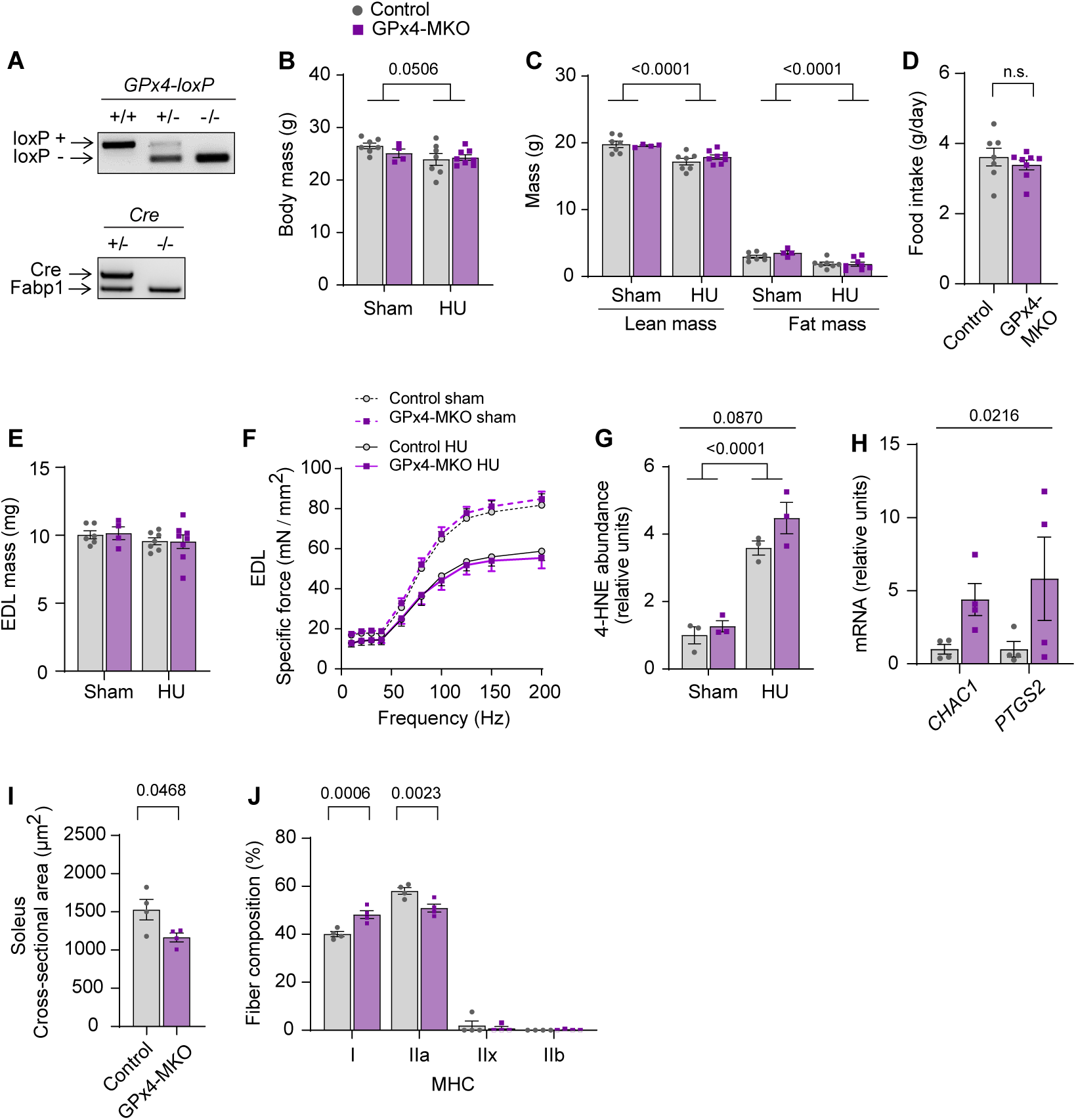
Additional data from GPx4-MKO mice. **(A)** Genotyping gel for GPx4cKO+/+ mice crossed with HSA-MerCreMer+/− mice to yield GPx4cKO+/+; HSA-MerCreMer+/− (GPx4-MKO) mice. Control littermates (GPx4cKO+/+; HSA- MerCreMer−/−) were used for experiments. **(B,C)** Body mass **(B)**, and body composition **(C)** (*n =* 7 for Ctrl sham, *n =* 4 for GPx4-MKO sham, *n =* 7 for Ctrl HU, *n =* 8 for GPx4- MKO HU). **(D)** Food intake during HU (*n =* 7 for Ctrl, *n =* 8 for GPx4-MKO). **(E,F)** EDL muscle mass (*n =* 6 for Ctrl sham, *n =* 4 for GPx4-MKO sham, *n =* 7 for Ctrl HU, *n =* 8 for GPx4-MKO HU) **(E)** and force-frequency curve from young mice (*n =* 6 for Ctrl sham, *n =* 4 for GPx4-MKO sham, *n =* 6 for Ctrl HU, *n =* 7 for GPx4-MKO HU) **(F)**. **(G)** Quantification of western blotting for 4-HNE from young mice (*n =* 3 per group). **(H)** mRNA levels of CHAC1 and PTGS2 in skeletal muscles (*n =* 4 per group). **(I,J)** Mean muscle fiber CSA (**i**) and fiber type composition **(J)** for soleus muscles following HU (*n =* 4 per group). Data are shown as the mean ± SEM. Statistical analyses in **(D)** and **(I)** were performed with an unpaired two-tailed t-test. Statistical analyses in **(B,C,E,F,G,H)** and **(J)** were performed with a two-way ANOVA and Tukey’s multiple comparison test.

**Supplemental Fig. S5.**
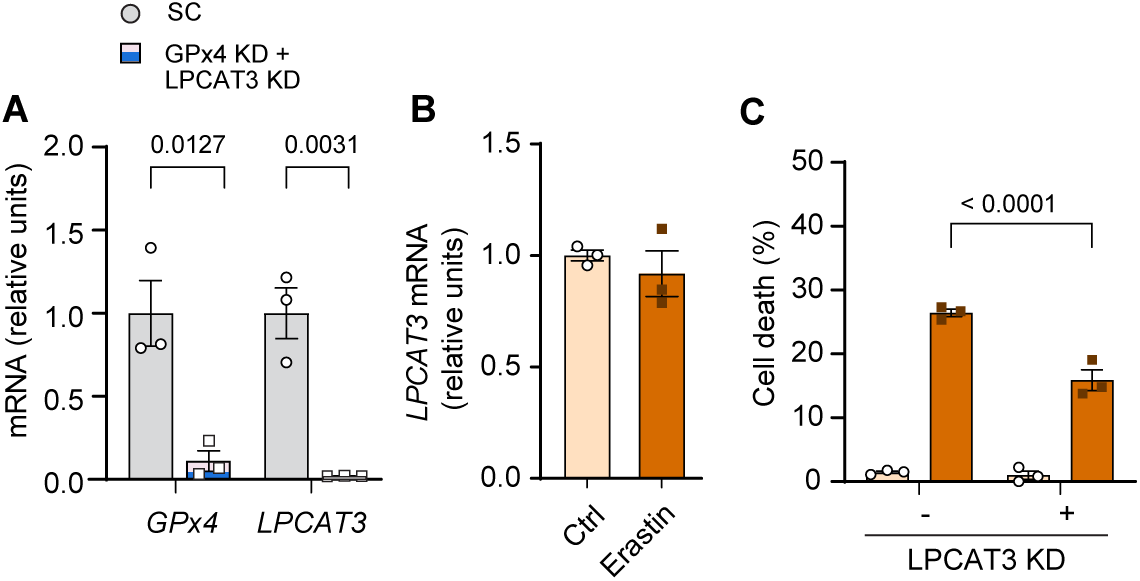
Additional data from LPCAT3KD. **(A)** mRNA levels of GPx4 and LPCAT3 in C2C12 myotubes with or without GPx4 and LPCAT3 double KD (*n =* 3 per group). **(B)** mRNA levels of LPCAT3 from erastin-stimulated myotubes (*n =* 3 per group). **(C)** Cell death levels from C2C12 myotubes with or without LPCAT3 KD and/or erastin (*n =* 3 independent repeats). Data are shown as the mean ± SEM. Statistical analyses in **(A)** and **(B)** were performed with an unpaired two-tailed t-test. Statistical analyses in **(C)** were performed with a two-way ANOVA and Tukey’s multiple comparison test.

**Supplemental Fig. S6.**
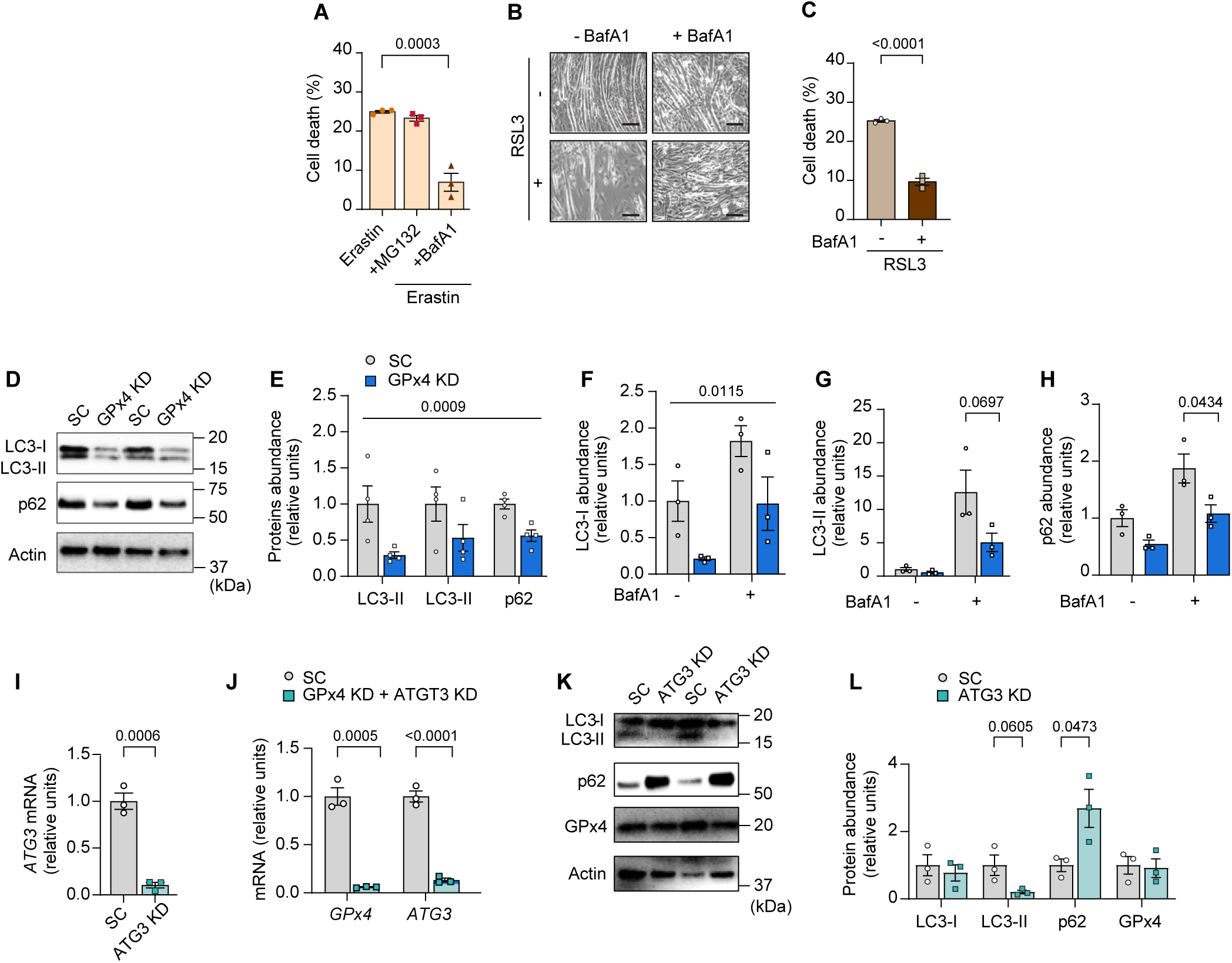
Autophagy-lysosome axis in LOOH-induced myotube atrophy. **(A)** Cell death levels in erastin-stimulated myotubes with MG132 or BaFA1 (*n =* 3 independent repeats). **(B,C)** Representative images **(B)** and cell death level **(C)** in RSL3-stimulated myotubes with BaFA1 (*n =* 3 independent repeats). **(D,E)** Immunoblotting **(D)** and quantification **(E)** of LC3-I, LC3-II, p62 (all static measurements without BafA1), and actin protein from C2C12 myotubes with or without GPX4 KD (*n =* 4 per group). **(F,G,H)** quantification of western blotting for LC3-I **(F)**, LC3-II **(G)**, and p62 **(H)** protein from GPx4 KD myotubes with or without BaFA1 (*n =* 3 per group). **(I)** mRNA levels of ATG3 in C2C12 myotubes with or without (*n =* 3 per group). **(J)** mRNA levels of GPx4 and ATG3 from C2C12 myotubes with double knockdown for ATG3 and GPx4 (*n =* 3 per group). **(K,L)** Immunoblotting **(K)** and quantification **(L)** of LC3, p62, and GPx4 protein from C2C12 myotubes with or without ATG3 KD (*n =* 3 per group). Data are shown as the mean ± SEM. Statistical analyses in **(C,I)** and **(L)** were performed with an unpaired two-tailed t-test. Statistical analyses in **(A)** were performed with a one-way ANOVA and Tukey’s multiple comparison test. Statistical analyses in **(E,F,G,H,I)**, and **(J)** were performed with a two-way ANOVA and Tukey’s multiple comparison test.

**Supplemental Fig. S7.**
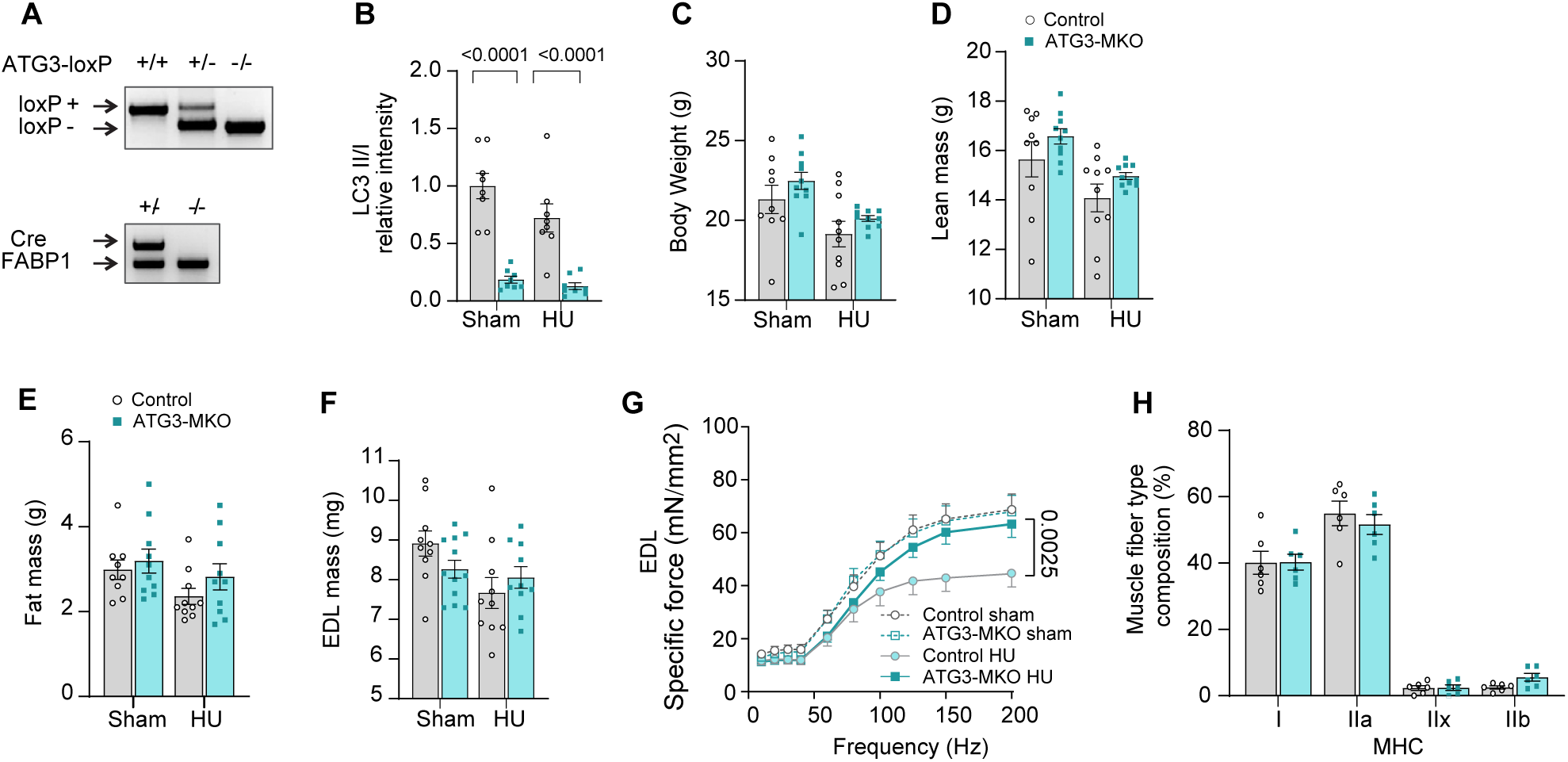
Additional data from ATG3-MKO mice. (**A**) Genotyping of ATG3cKO+/+ mice crossed with HSA-MerCreMer+/− mice to yield ATG3cKO+/+; HSA- MerCreMer+/− (ATG3-MKO) mice. Control littermates (ATG3cKO+/+; HSA-MerCreMer−/−) were used for experiments. **(B)** Quantification of LC3-II/I ratio in muscles from control and ATG3-MKO mice (*n =* 8 per group). **(C,D,E)** Body weight **(C)**, lean mass **(D)**, and fat mass **(E)** from control and ATG3-MKO mice (control sham *n =* 9, ATG3-MKO sham *n =* 10, control HU *n =* 10, ATG3-MKO HU *n =* 10). (**F**) EDL muscle mass (control sham *n =* 10, ATG3-MKO sham *n =* 12, control HU *n =* 10, ATG3-MKO HU *n =* 10). (**G**) Force frequency curve for EDL muscle (control sham *n =* 9, ATG3-MKO sham *n =* 8, control HU *n =* 10, ATG3-MKO HU *n =* 9). (**H**) Fiber type composition for SOL muscle (control HU *n =* 6, ATG3MKO HU *n =* 6). Data are shown as the mean ± SEM. Statistical analyses in **(B,C,D,E,F,G)** and **(H)** were performed with a two-way ANOVA and Tukey’s multiple comparison test.

**Supplemental Fig. S8.**
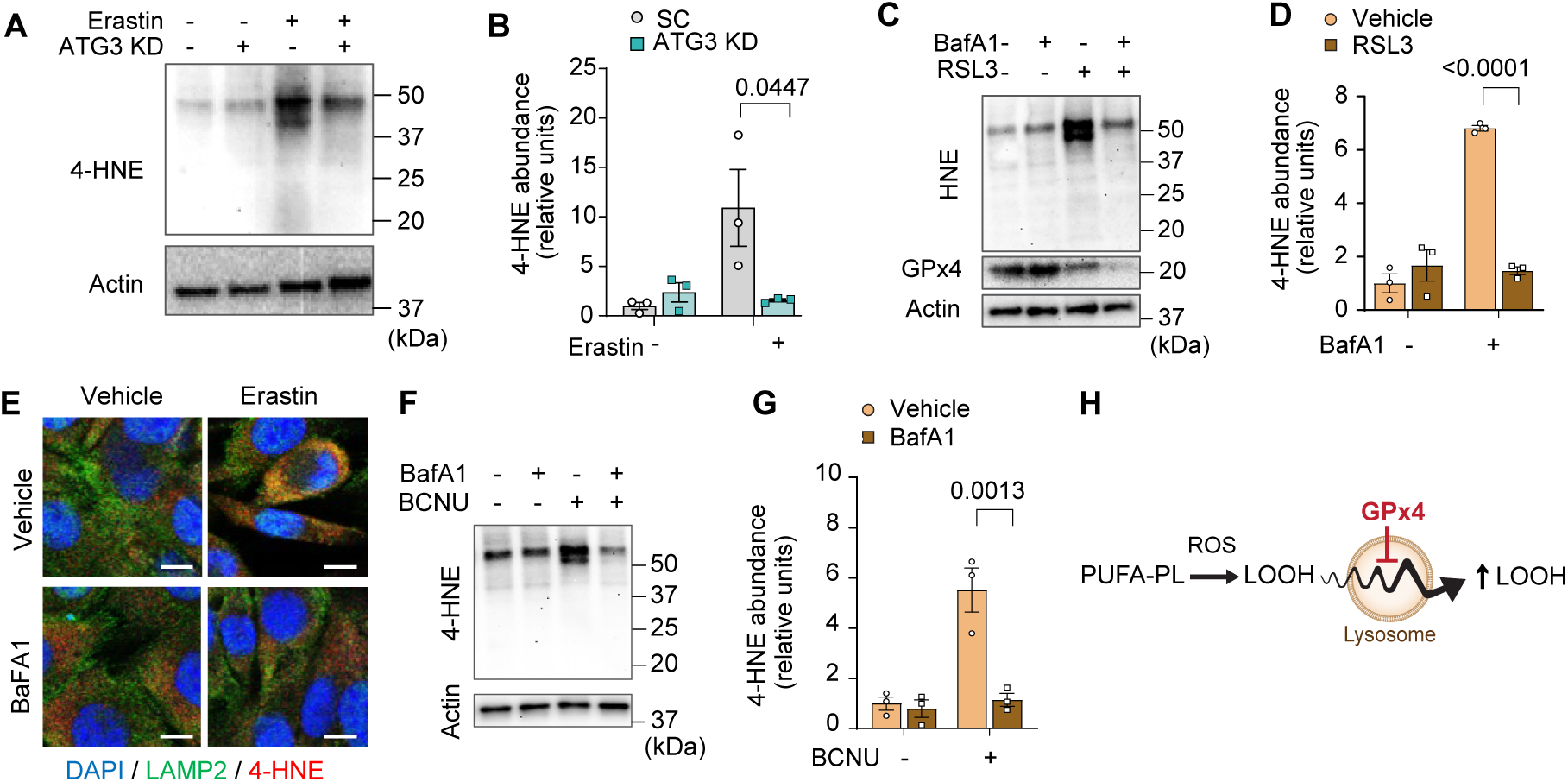
Inhibition of autophagy-lysosome axis prevents accumulation of LOOH. **(A,B)** Immunoblotting **(A)** and quantification **(B)** of 4-HNE protein from C2C12 myotubes with or without erastin and/or ATG3 KD (*n =* 3 per group). **(C,D)** Immunoblotting **(C)** and quantification **(D)** of 4-HNE protein from C2C12 myotubes with or without erastin and/or BaFA1 (*n =* 3 per group). **(E)** Confocal immunofluorescence of LAMP2 (lysosome) and 4-HNE in erastin-stimulated C2C12 myotubes with or without BaFA1. Scale bar, 10µm. **(F,G)** Immunoblotting **(F)** and quantification **(G)** of 4-HNE protein from BCNU-stimulated C2C12 myotubes with or without BafA1. **(H)** A novel role of lysosome in amplifying LOOH. Data are shown as the mean ± SEM. Statistical analyses in **(B,D)** and **(G)** were performed with a two-way ANOVA and Tukey’s multiple comparison test.

**Supplemental Fig. S9.**
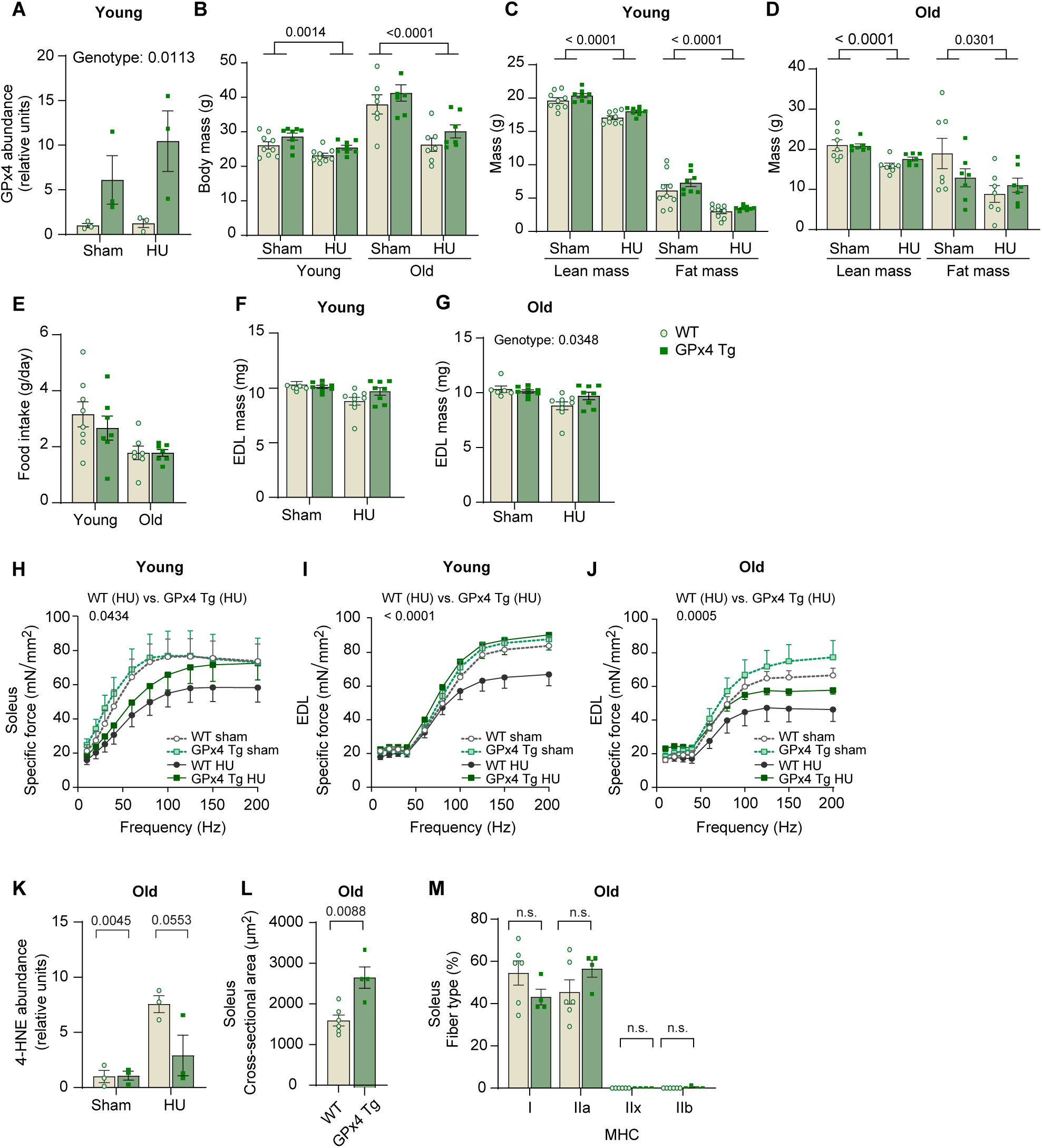
Additional data from GPx4Tg mice. (**A**) Quantification of GPx4 proteins from WT or GPx4Tg with or without HU (*n =* 3 per group). **(B)** Body mass from young (*n =* 9 for WT, *n =* 8 for GPx4Tg) and old (*n =* 7 for WT, *n =* 7 for GPx4Tg) mice. **(C,D)** body composition from young mice **(C)** and old mice **(D)**. **(E)** Food intake during HU from young and old mice. **(F,G)** EDL muscle mass from young (*n =* 6 for WT sham, *n =* 8 for GPx4Tg sham, *n =* 9 for WT HU, *n =* 8 for GPx4Tg HU) **(F)** and old (*n =* 7 per group) **(G)** mice. **(H)** Force-frequency curve in soleus muscle from young mice (*n =* 5 for WT sham, *n =* 6 for GPx4Tg sham, *n =* 9 for WT HU, *n =* 8 for GPx4Tg HU). **(I,J)** Force-frequency curve in EDL muscle from young mice (*n =* 5 for WT sham, *n =* 7 for GPx4Tg sham, *n =* 9 for WT HU, *n =* 8 for GPx4Tg HU) **(I)** and old (*n =* 5 for WT sham, *n =* 5 for GPx4Tg sham, *n =* 5 for WT HU, *n =* 6 for GPx4Tg HU) **(J)** mice. **(K)** Quantification of muscle 4-HNE proteins from old mice (*n =* 3 per group). **(L,M)** Mean muscle fiber CSA **(L)** and fiber type composition **(M)** for soleus muscles from old mice following HU (*n =* 6 for WT, *n =* 4 for GPx4Tg). Data are shown as the mean ± SEM. Statistical analyses in **(L)** were performed with an unpaired two-tailed t-test. Statistical analyses in **(A-M)** were performed with a two-way ANOVA and Tukey’s multiple comparison test.

**Supplemental Fig. S10.**
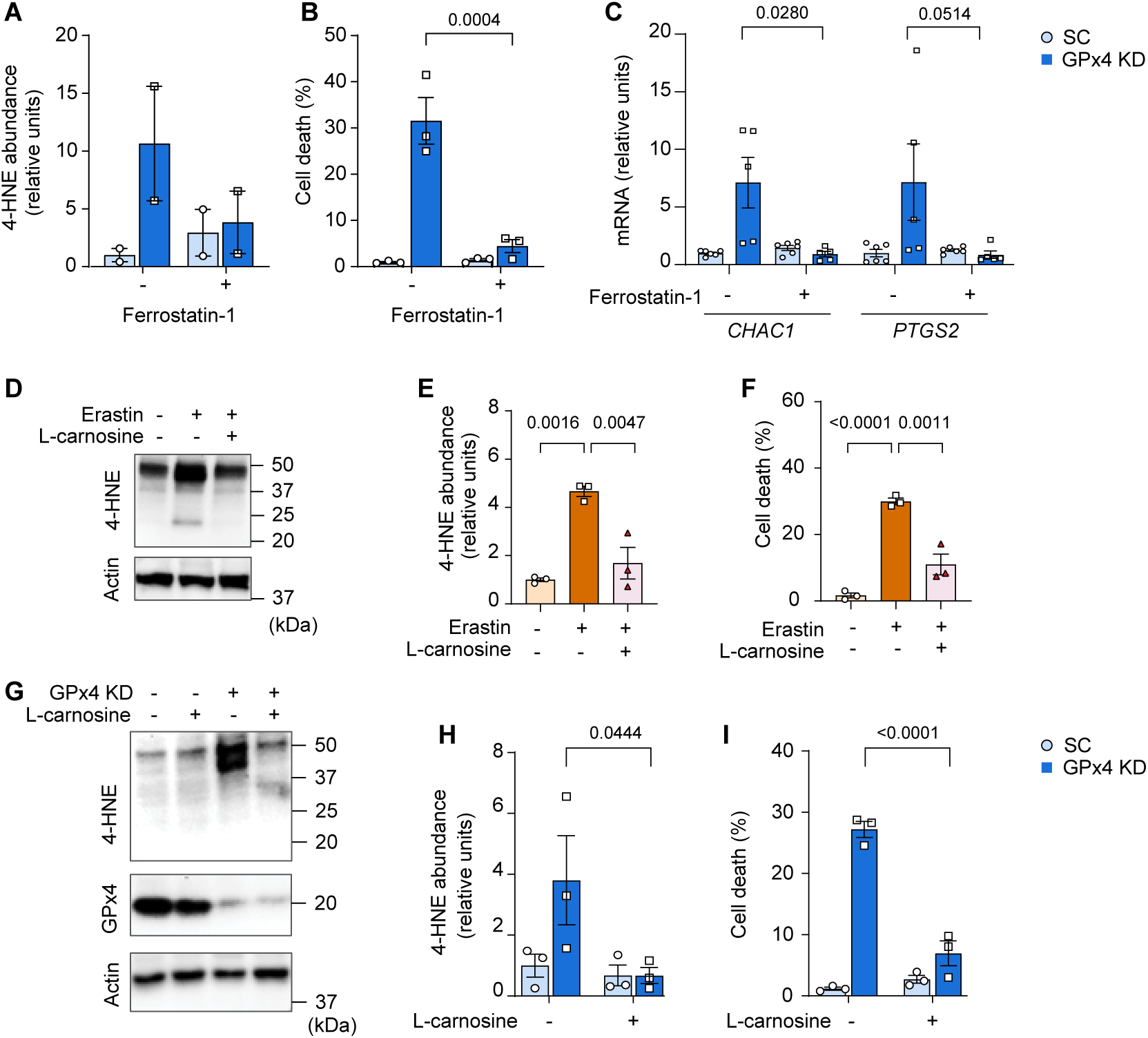
Effects of ferrostatin-1 or L-carnosine treatment in vitro. **(A,B,C)** Quantification of 4-HNE proteins (*n =* 2 per group) **(A)**, cell death level (*n =* 3 independent repeats) **(B)**, and mRNA levels of CHAC1 and PTGS2 with or without GPx4 KD and/or Ferrostatin-1 (*n =* 6 for SC, *n =* 5 for GPx4 KD) **(C)** in C2C12 myotubes. **(D,E,F)** Immunoblotting **(D)** and quantification **(E)** of 4-HNE, and actin (*n =* 3 per group) and cell death level (*n =* 3 independent repeats) **(F)** with or without erastin and/or L-carnosine in C2C12 myotubes. **(G,H,I)** Immunoblotting **(G)** and quantification **(H)** of 4-HNE, and actin (*n =* 3 per group), and **(I)** cell death level (*n =* 3 independent repeats) in C2C12 myotubes with or without GPx4 KD and/or L-carnosine. Data are shown as the mean ± SEM. Statistical analyses in **(E)**, and **(F)** were performed with a one-way ANOVA and Tukey’s multiple comparison test. Statistical analyses in **(A,B,C,H)** and **(I)** were performed with a two-way ANOVA and Tukey’s multiple comparison test.

**Supplemental Fig. S11.**
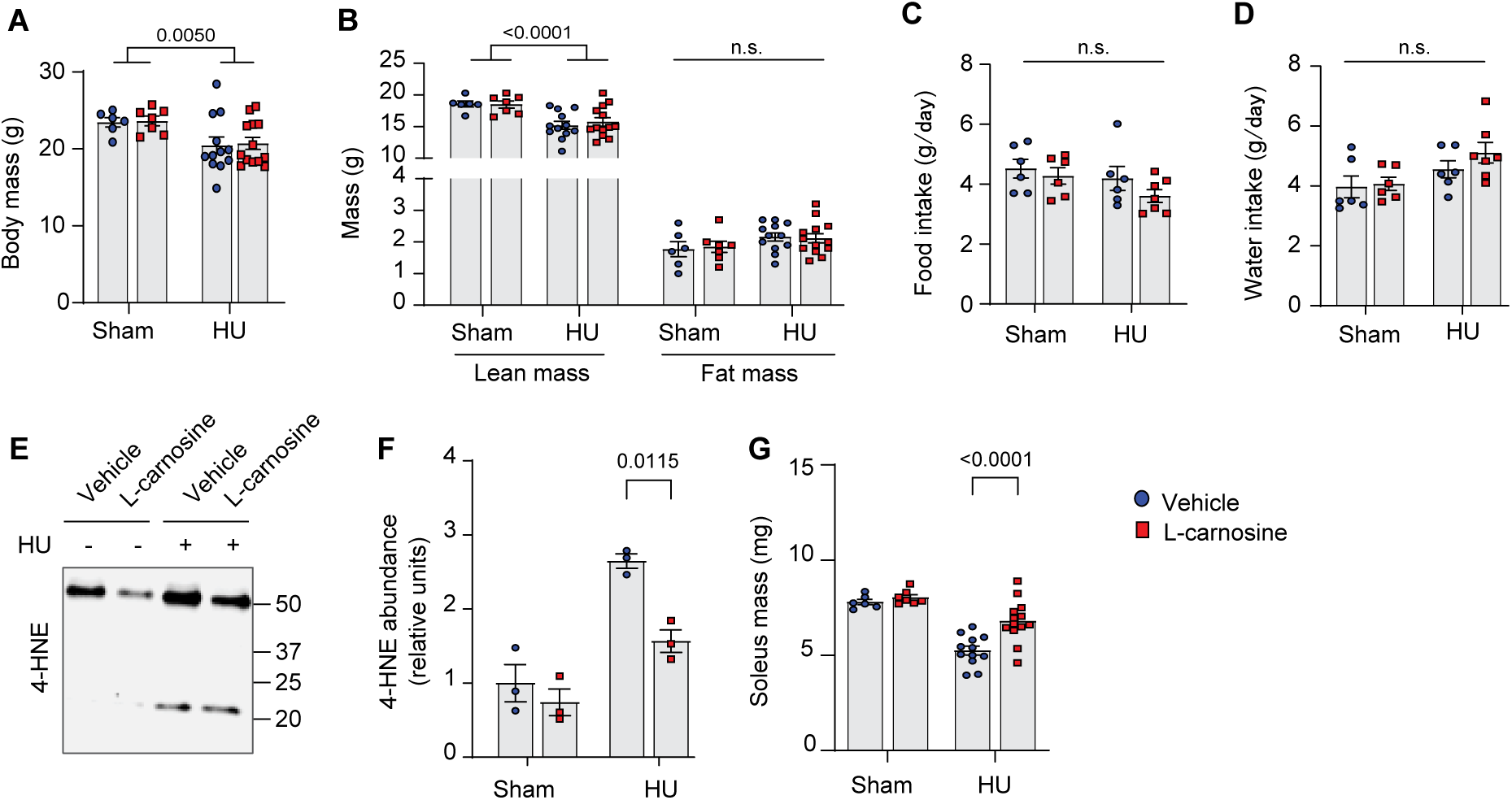
L-carnosine treatment partly ameliorates disuse-induced muscle atrophy. **(A,B,C,D,E,F,G)** Body mass **(A)**, body composition (*n =* 6 for vehicle sham, *n =* 7 for L-carnosine sham, *n =* 12 for vehicle HU, *n =* 13 for L-carnosine HU) **(B)**, food intake **(C)**, water intake (*n =* 6 for vehicle sham, *n =* 6 for L-carnosine sham, *n =* 6 for vehicle HU, *n =* 7 for L-carnosine HU) (**D**), immunoblotting **(E)** and quantification **(F)** of 4-HNE (*n =* 3 per group), and soleus muscle mass (*n =* 6 for vehicle sham, *n =* 7 for L-carnosine sham, *n =* 12 for vehicle HU, *n =* 13 for L-carnosine HU) **(G)**. Data are shown as the mean ± SEM. Statistical analyses in **(A,B,C,D,F)** and **(G)** were performed with a two-way ANOVA and Tukey’s multiple comparison test.

**Supplemental Fig. S12.**
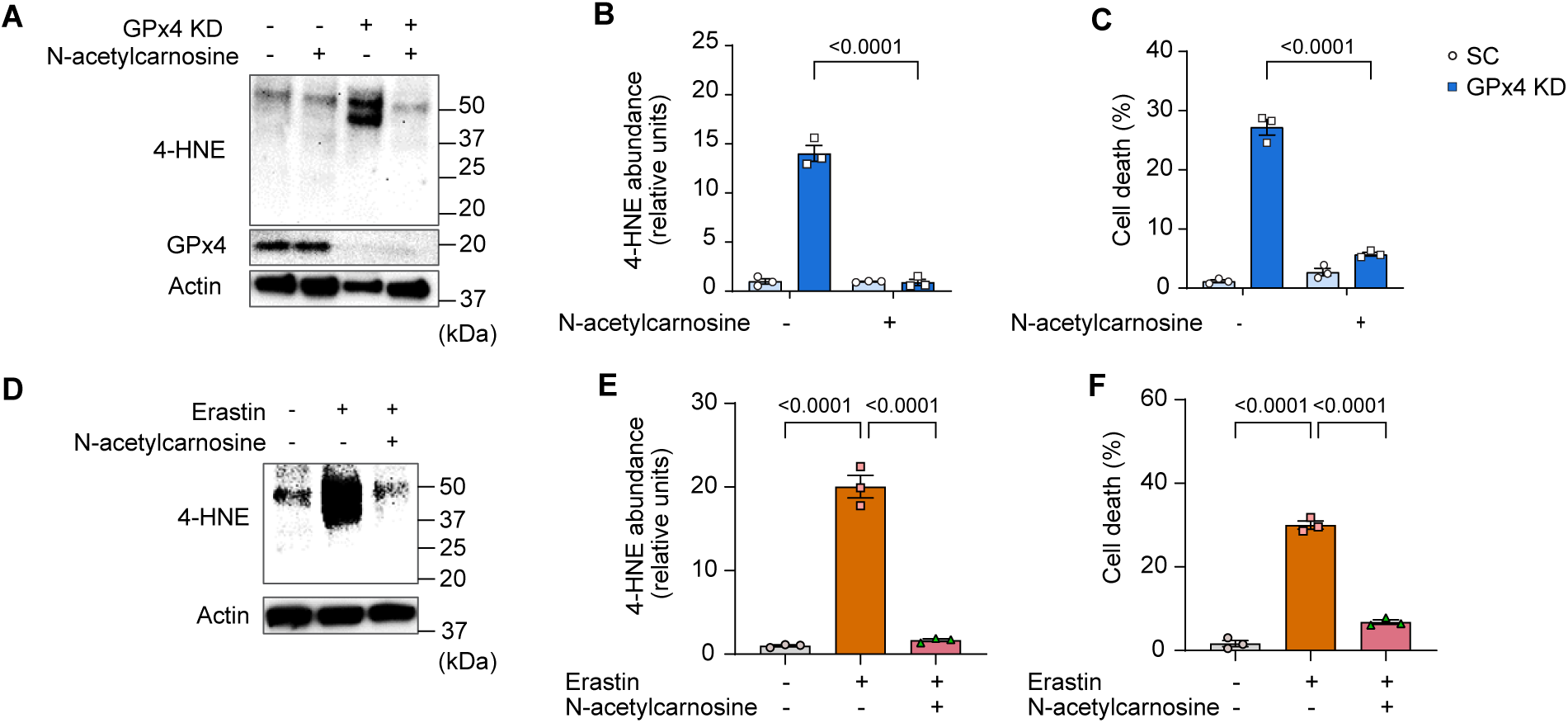
Effects of N-acetylcarnosine treatment in vitro. **(A,B,C)** Immunoblotting **(A)** and quantification **(B)** for 4-HNE, GPx4 and actin (*n =* 3 per group) and cell death level (*n =* 3 independent repeats) **(C)** from C2C12 myotubes with or without GPx4 KD and/or N-acetylcarnosine. **(D,E,F)** Immunoblotting **(D)** and quantification **(E)** of 4-HNE and actin (*n =* 3 per group) and cell death level (*n =* 3 independent repeats) **(F)** in C2C12 myotubes with or without erastin and/or N-acetylcarnosine. Data are shown as the mean ± SEM. Statistical analyses in **(B)** and **(C)** were performed with a two-way ANOVA and Tukey’s multiple comparison test. Statistical analyses in **(E)** and **(F)** were performed with a one-way ANOVA and Tukey’s multiple comparison test.

**Supplemental Fig. S13.**
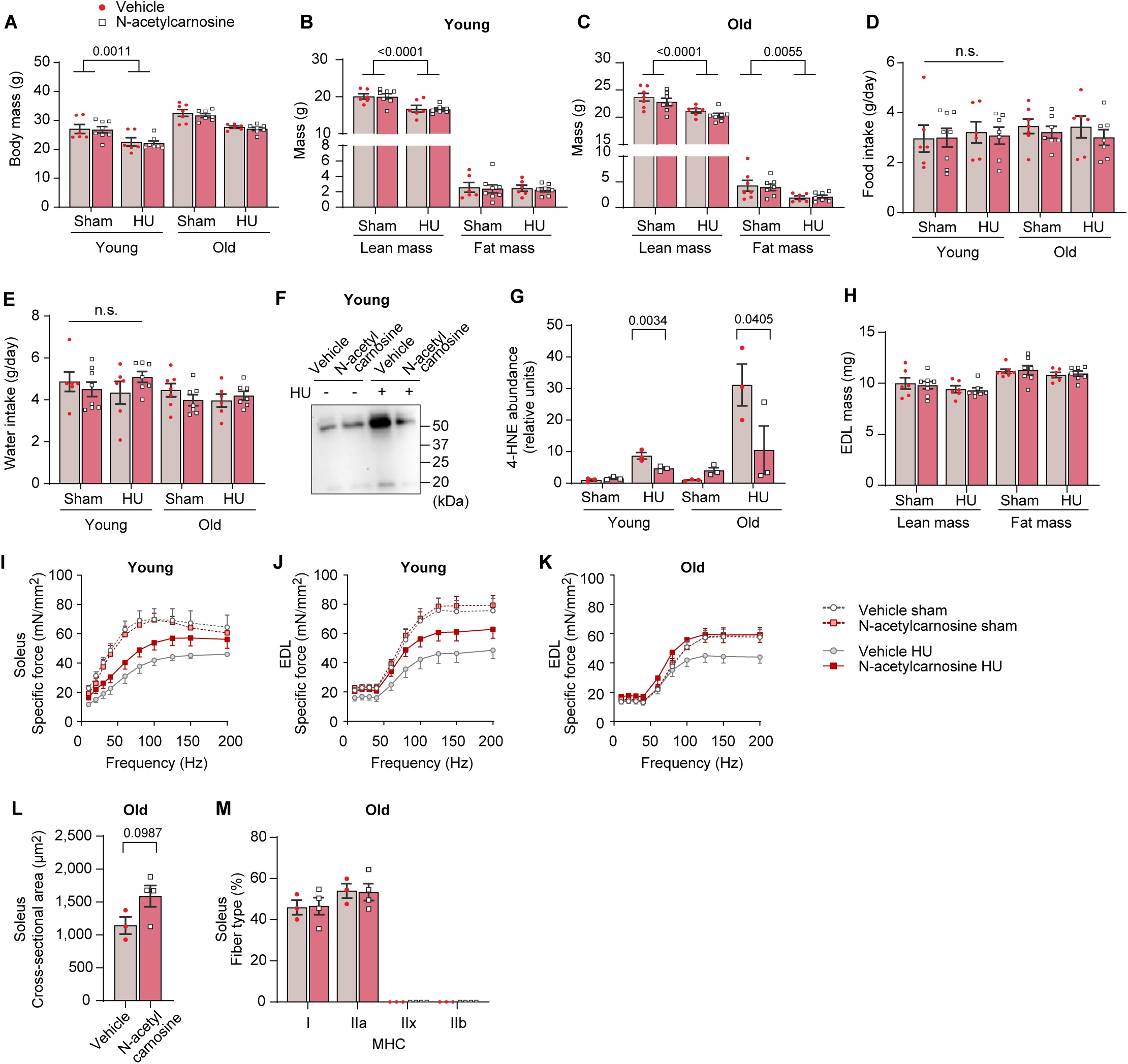
Additional data from N-acetylcarnosine treatment in vivo. **(A,B,C)** Body mass **(A)**, body composition from young mice **(B)** and old mice **(C)**. **(D,E,F,G)** Food intake **(D)**, water intake **(E)**, immunoblotting **(F)** and quantification **(G)** of 4-HNE (*n =* 3 per group). **(H)** EDL muscle mass from young mice and old mice. **(I)** Force-frequency curve in soleus muscle from young mice (*n =* 5 for vehicle sham, *n =* 6 for N-acetylcarnosine sham, *n =* 5 for vehicle HU, *n =* 5 for N-acetylcarnosine HU). **(J,K)** Force-frequency curve in EDL muscle from young mice (*n =* 7 for vehicle sham, *n =* 7 for N-acetylcarnosine sham, *n =* 6 for vehicle HU, *n =* 7 for N-acetylcarnosine HU) **(J)** and old mice (*n =* 6 for vehicle sham, *n =* 7 for N-acetylcarnosine sham, *n =* 6 for vehicle HU, *n =* 6 for N-acetylcarnosine HU) **(K)**. **(L,M)** Average of muscle fiber CSA (**l**) and fiber type composition **(M)** for soleus muscles from old mice following HU (*n =* 3 for vehicle, *n =* 4 for N-acetylcarnosine). Data are shown as the mean ± SEM. Statistical analyses in **(L)** were performed with an unpaired two-tailed t-test. Statistical analyses in **(A,B,C,D,E,G,H,I,J,K)** and **(M)** were performed with a two-way ANOVA and Tukey’s multiple comparison test.

